# PINK1-dependent phosphorylation of Serine111 within the SF3 motif of Rab GTPases impairs effector interactions and LRRK2 mediated phosphorylation at Threonine72

**DOI:** 10.1101/764019

**Authors:** Sophie Vieweg, Katie Mulholland, Bastian Bräuning, Nitin Kachariya, Yu-Chiang Lai, Rachel Toth, Michael Sattler, Michael Groll, Aymelt Itzen, Miratul M. K. Muqit

## Abstract

Loss of function mutations in the PINK1 kinase are causal for autosomal recessive Parkinson disease (PD) whilst gain of function mutations in the LRRK2 kinase cause autosomal dominant PD. PINK1 indirectly regulates the phosphorylation of a subset of Rab GTPases at a conserved Serine111 (Ser111) residue within the SF3 motif. Using genetic code expansion technologies we have produced stoichiometric Ser111-phosphorylated Rab8A revealing impaired interactions with its cognate guanine nucleotide exchange factor (GEF) and GTPase activating protein (GAP). In a screen for Rab8A kinases we identify TAK1 and MST3 kinases that can efficiently phosphorylate the Switch II residue Threonine72 (Thr72) in a similar manner as LRRK2. Strikingly we demonstrate that Ser111 phosphorylation negatively regulates the ability of LRRK2 but not MST3 or TAK1 to phosphorylate Thr72 *in vitro* and demonstrate an interplay of PINK1- and LRRK2-mediated phosphorylation of Rab8A in cells. Finally, we present the crystal structure of Ser111-phosphorylated Rab8A and NMR structure of Ser111-phosphorylated Rab1B that does not demonstrate any major changes suggesting that the phosphorylated SF3 motif may disrupt effector-Switch II interactions. Overall, we demonstrate antagonistic regulation between PINK1-dependent Ser111 phosphorylation and LRRK2-mediated Thr72 phosphorylation of Rab8A suggesting that small molecule activators of PINK1 may have therapeutic potential in patients harbouring LRRK2 mutations.

## Introduction

Autosomal recessive mutations in PTEN-induced kinase 1 (PINK1) represent the second most frequent cause of early-onset Parkinson’s disease [1]. PINK1 encodes a protein kinase that is localised to the mitochondria via a N-terminal mitochondrial targeting domain. In response to chemical uncouplers (e.g. carbonyl cyanide m-chlorophenyl hydrazone (CCCP)) that induce mitochondrial depolarisation, PINK1 is stabilised and activated and directly phosphorylates Parkin at Serine 65 (Ser65) within its N-terminal Ubiquitin-like (Ubl) domain [2, 3] as well as the equivalent Ser65 residue of ubiquitin [4–7]. The phosphorylation of both of these residues is required for maximal activation of Parkin E3 ligase activity that induces ubiquitylation of a myriad of substrates and acts as a signal for the elimination of damaged mitochondria via mitophagy [4–7].

Previous work has shown that upon activation, PINK1 indirectly induces the phosphorylation of a subset of Rab GTPases including Rab 1B, 8A, 8B, and 13 at a highly conserved Serine residue (Ser111 in Rab8A), that lies within the RabSF3 motif [8]. Of relevance to Parkinson’s disease mechanisms, loss of function mutations of PINK1 abolish Rab protein phosphorylation as assessed in primary mouse embryonic fibroblasts (MEFs) from PINK1 knockout mice or human adult fibroblasts derived from patients harbouring homozygous PINK1 Q456X mutations [8].

Recent studies have also linked other Parkinson’s genes to the regulation of Rab proteins. Autosomal dominant activating mutations of LRRK2 represent the commonest genetic cause of familial PD accounting for ∼5% of all cases and clinical LRRK2 inhibitor trials have recently started (Denali Therapeutics). LRRK2 phosphorylates a highly conserved Threonine residue within the Switch II domain of 14 Rab GTPase isoforms (Thr72 in Rab8A) that lies upstream of the PINK1-regulated Ser111 site [9, 10]. Furthermore, autosomal dominant missense mutations of the vacuolar protein sorting-associated protein VPS35 (D620N) are associated with a substantial elevation of LRRK2-mediated Rab phosphorylation [11].

The identity of the upstream Rab Ser111 kinase remains unknown and this has hampered progress to determine the impact of PINK1-dependent Ser111 phosphorylation on the function of Rab GTPases. Another major unanswered question is whether PINK1 lies in signalling networks with other PD gene pathways. Herein we undertake a screen to identify Rab protein kinases and discover that TAK1 and MST3 can efficiently phosphorylate Thr72 equivalent to the LRRK2 site. We have also employed genetic code expansion technologies to produce milligram quantities of Ser111-phosphorylated Rab8A and Rab1B, that has enabled investigation of how phosphorylation affects Rab function and has revealed an antagonistic role between PINK1 and LRRK2-mediated phosphorylation of Rabs.

## Materials and Methods

### Reagents

[γ-^32^P] ATP was from PerkinElmer, MLi-2 was obtained from Merck [30]. LRRK2 (G2019S) Recombinant Human Protein (residues 970-2527 #10499963) was purchased from Invitrogen and all additional kinases used in the *in vitro* screen were from MRC PPU Reagents and Services – including TAK1 (1-303), TAB1 (437-504) fusion DU753 and MST3 (1-431) DU number: 30889 N-terminal GST. All mutagenesis was carried out using the QuikChange site-directed mutagenesis method (Stratagene) with KOD polymerase (Novagen). All DNA constructs were verified by DNA sequencing, which was performed by The Sequencing Service, School of Life Sciences, University of Dundee, using DYEnamic ET terminator chemistry (Amersham Biosciences) on Applied Biosystems automated DNA sequencers. DNA for bacterial protein expression was transformed into *E. coli* BL21 DE3 RIL (codon plus) cells (Stratagene). All cDNA plasmids, antibodies and recombinant proteins generated for this study are available to request through our reagents website https://mrcppureagents.dundee.ac.uk/ or from the Itzen laboratory.

### Expression of non-phosphorylated (WT) Rab proteins

WT-Rab1B(3-174) was expressed in fusion with a N-terminal His_6_-MBP tag (pMAL) or a C-terminal His_6_ tag (pNHD), while WT-Rab8A(6-176) was fused to a N-terminal His_6_ tag or a C-terminal His_6_ tag (pET19). The N-terminal Rab1B and Rab8A fusion constructs contained a TEV cleavage site to remove the His_6_-MBP tag or His_6_ tag from the N-terminus of the Rab proteins. The C-terminal His_6_ tags remained on the Rab proteins. All WT-Rab proteins were expressed in BL21(DE3) *E.coli* in LB medium at 25°C overnight, and their expression was induced with 0.5 mM IPTG at OD_600nm_ = 0.8. In contrast to the Rab1B proteins, the Rab8A proteins were co-expressed with the chaperone GroEL/S (pGro7, Takara). The expression of the GroEL/S chaperone was auto-induced by supplementing the LB-medium with 1 mg/mL arabinose.

### Phosphorylation of Rab proteins by genetic code expansion

For the incorporation of phospho-serine (pSer) during the expression of Rab1B (3-174) and Rab8A (6-176) proteins, we have employed a genetic code expansion strategy published by Rogerson and colleagues [15]. For this purpose, we introduced an amber stop codon at the amino acid position Ser111 via site-specific mutagenesis. We co-transformed the obtained constructs, Rab1B(S111(TAG))-His_6_ (pNHD) and Rab8A(S111(TAG))-His_6_ (pNHD), together with the pKW2 plasmid encoding for the orthogonal tRNA/tRNA-synthetase pair (SepRS(2)/pSer tRNA(B4)CUA) and the optimized elongation factor EF-Sep into the ΔserB-BL21(DE3) *E.coli* expression strain. In the case of pSer111-Rab8A, a third plasmid encoding for the GroEL/S chaperone (pET19) was transformed into the expression strain. The cells were grown in TB medium at 37°C and 180 rpm. The culture was supplemented with 2 mM O-Phospho-L-serine (Sigma) at OD_600nm_ = 0.3. The expression of all plasmids was induced simultaneously by the addition of 1 mM IPTG at OD_600nm_ = 0.8-0.9. The expression was carried out overnight at 30°C for pSer111-Rab1B and at 25°C for pSer111-Rab8A.

### Purification of Rab proteins

Cells were harvested and resuspended in buffer A (50 mM HEPES, 500 mM LiCl, 10 mM imidazole, 1 mM MgCl2, 10 µM GDP, 2 mM β-mercaptoethanol, pH 8.0) containing 1 mM PMSF and DNaseI. Cells were lysed on ice by sonication (60% amplitude, 5 min, pulse 5 s on and 15 s off), and insoluble cell debris was removed by centrifugation (48254.4 x g, 45 min, 4°C). All Rab proteins were purified from the supernatant by nickel affinity chromatography (5 ml HiTrap™ Chelating HP column, GE Healthcare). Non-specifically bound proteins were removed by a step wash with buffer B (50 mM HEPES, 500 mM LiCl, 500 mM imidazole 1 mM MgCl2, 10 µM GDP, 2 mM β-mercaptoethanol, pH 8.0): 5% buffer B to remove impurities from non-phosphorylated (WT) Rab proteins, or three consecutive washes with 3%, 5% and 7% buffer B to purify phosphorylated Rab proteins, respectively. WT-Rab proteins eluted with a linear gradient from 5-60% buffer B in 100 ml, and phosphorylated Rab proteins eluted with a linear gradient from 7-50% buffer B in 50 ml. The collected fractions were analyzed by SDS-PAGE. Fractions containing the desired His_6_-tagged protein were concentrated to 2 ml using Amicon® Ultra Centrifugal Filters (10,000 MWCO, Merck Millipore), and were directly injected into a size-exclusion chromatography column (Superdex 75 16/60, GE Healthcare) equilibrated with buffer C (20 mM HEPES, 50 mM NaCl, 1 mM MgCl2, 10 µM GDP, 1 mM DTT, pH 7.5). The purity of the fractions from the size-exclusion chromatography was determined by SDS-PAGE, and the clean fractions were pooled, concentrated using Amicon® Ultra Centrifugal Filters (10,000 MWCO, Merck Millipore), and frozen in liquid nitrogen. The protein identity and integrity was confirmed by LC-MS. Note that Rab proteins with a N-terminal His_6_ tag or His_6_-MBP tag were treated with TEV protease after the nickel affinity purification, and dialyzed against 5L of buffer D (20 mM HEPES, 100 mM NaCl, 10 µM GDP, 2 mM β-mercaptoethanol, pH 8.0) at 4°C overnight. The cleaved N-terminal tags were separated from the Rab proteins by a second nickel affinity purification prior to the size-exclusion chromatography.

### Nucleotide exchange of Rab proteins

The active form of Rab proteins was prepared by incubating the Rab proteins with 5 mM EDTA and 20 molar equivalents of GppNHp or GTP at 25°C for 3 h. The excess nucleotide was removed by buffer exchange using NAP5 columns (GE healthcare) or size-exclusion chromatography columns (Superdex 75 10/300, GE Healthcare) equilibrated with buffer E (20 mM HEPES, 50 mM NaCl, 1 mM MgCl2, 10 µM GTP or GppNHp, 1 mM DTT, pH 7.5). The nucleotide loading of the Rab proteins was determined by C18 RP-HPLC. Therefore, 50 µM of Rab protein were heat denatured at 95°C for 5 min and spun down at maximum speed. The supernatant was injected into a C18 column equilibrated with 50 mM KPi (pH 6.6), 10 mM tetra-n-butylammonium bromide and 17% ACN. A mixture of guanosine nucleotides (GMP, GDP and GTP) and pure GppNHp were used as standards to determine the nucleotide loading state of the Rab proteins.

### Guanosine exchange factor (GEF) and GTPase activating protein (GAP) activity assay

GDP- or GTP-bound Rab proteins were diluted in buffer F (20 mM HEPES, 50 mM NaCl, 1 mM MgCl_2_, pH 7.5) to a concentration of 4 µM, and transferred into a Quartz SUPRASIL cuvette (Hellma Analytics, Germany). The intrinsic tryptophane (Trp) fluorescence was recorded at 348 nm (ex = 297 nm) at 25°C with a Fluoromax-4 fluorescence spectrometer (HORIBA Jobin Yvon). When the Trp fluorescence signal stabilized, 50 µM of GTP were added into the cuvette containing 4 µM of Rab:GDP followed by the addition of a GEF enzyme (0.04 µM DrrA or 2 µM Rabin8) to record the nucleotide exchange reaction (GDP→GTP). To monitor the GTP hydrolysis reaction, 0.04 µM of the GAP TBC1D20 were mixed into 4 µM Rab:GTP. Based on the decrease or increase in Trp fluorescence over time, the GDP-GTP exchange rates and GTP hydrolysis rates (k_obs_) were calculated for each Rab protein using the following exponential equation: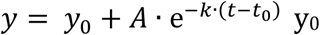: final fluorescence value (end point) A: amplitude of fluorescence change t: time in seconds t_0_: initial time point of the reaction

### Isothermal titration calorimetry (ITC)

Rab8A-Mical-1 binding studies were conducted with a MicroCal PEAQ-ITC device (Malvern Instruments Ltd, UK). WT-Rab8A-His_6_ and pSer111-Rab8A-His_6_ were loaded with GppNHp as described above, and afterwards dialyzed together with Mical-1 against 5L of buffer G (20 mM HEPES, 50 mM NaCl, 2 mM MgCl_2_, 1 mM TCEP, pH 7.5) at 4°C overnight. To determine the dissociation constant (K_D_), 400 µM of WT-Rab8A-His_6_: GppNHp or pSer111-Rab8A-His_6_:GppNHp were titrated into the cell containing 40 µM of Mical-1. We performed 26 injections of 1.5 µl with a spacing of 120 s at 500 rpm stirring speed at 25°C. The data analysis was carried out with the integrated MicorCal software, and the raw data was fitted to a one-site binding model. For every experiment one technical replicate was recorded.

### Thermal Shift Assay (TSA)

The thermal stability or melting temperatures (Tm) of the Rab proteins was determined using a SYPRO® orange-based TSA. Therefore, 2 µg of Rab protein were mixed with 5x SYPRO® orange protein gel stain (Sigma) in a volume of 20 µl. The TSA assay was performed in a CFX Connect™ real-time system (BioRad) using a stepwise temperature increase from 25°C to 95° in 1°C per min (71 cycles). The excitation wavelength was set to 470 nm and emission was detected at 570 nm. The raw data was plotted (fluorescence F vs. temperature T) in Excel, and the Tm values (inflection points) were derived from the second derivate (*f*′′(*T*) = 0) of each data set.

### Liquid chromatography mass spectrometry (LC-MS)

Protein samples (0.1 mg/ml, 1 µl) were injected with an UltiMate3000® HPLC system (UHPLC^+^ focused, Dionex) into a ProSwift™ RP-4H column (1x 50 mm, Thermo) at a flow rate of 0.7 ml/min. The proteins eluted with a linear gradient of 5-100% acetonitrile (0.1% formic acid) in 6 min. The desalted samples were ionized and analyzed by a LCQ fleet^™^ system (Thermo) combining electro spray ionization (ESI) with an ion trap mass analyzer.

### Phos-tag™ acrylamide gel analysis

SDS-PAGE gels (12% acrylamide) were prepared with 50 µM Phos-tag™ Acrylamide AAL-107, a phosphate-binding compound from Wako, and 100 µM MnCl_2_. 500 ng of the phosphorylated Rab1B proteins (GDP/GppNHp) were loaded on the gel, together with the non-phosphorylated Rab1B proteins as negative controls.

### Kinase assays

LRRK2 kinase assays were based on previous assay [10]. Assays were performed with a final volume of 25 µl with 2 µg Rab8A substrate, 100 ng LRRK2 G2019S (PV4881 Fisher Scientific), in 50 mM Tris-HCl (pH 7.5), 10 mM MgCl_2_ and 0.1 mM [γ-^32^P] ATP (∼300–500 c.p.m./pmol, PerkinElmer) in the absence or presence of MLi-2 (1 µM). Assays were incubated for 30 minutes at 30 ◦C, with shaking at 1050 r.p.m. and quenched through addition of SDS sample loading buffer with 1 % beta-mercaptoethanol and heating at 75 ◦C for 10 minutes. Reaction mixtures were resolved by SDS-PAGE. Proteins were detected by Coomassie staining and gels imaged with an Epson scanner, prior to gel drying (Bio-Rad). Incorporation of [γ-^32^P] ATP into substrates was analysed by autoradiography, using Amersham hyperfilm. Quantification of [γ - ^32^P] ATP into substrates was performed by Cerenkov counting of SDS-PAGE gel bands.

### Crystallization of pSer111-Rab8A:GDP/GppNHp

Crystallization trials of human pSer111-Rab8A:GDP were performed by the sitting-drop vapor diffusion method, using protein concentrations of 16 mg/mL and a protein:reservoir drop ratio of 1:1. Suitably sized crystals were grown in 0.2 M magnesium acetate, 0.1 M sodium cacodylate pH 6.5, 20% PEG 8000. Crystals were cryoprotected in reservoir solution supplemented with 20% glycerol prior to flash freezing in liquid nitrogen. Initial sparse-matrix screening of conditions yielded several hits for pSer111-Rab8A:GppNHp, which failed to produce well diffracting crystals despite fine-screening efforts. Using numerous crystals grown in 0.1M MES pH 6.5 and 25% PEG 6000/8000, and in 0.1M Hepes pH 7.5 and 25% PEG 3000/4000/6000 as seeds, the method of random microseed matrix seeding [31] was used to produce well diffracting crystals grown from a protein concentration of 11 mg/mL and a reservoir composed of 0.1 M Tris-HCl pH 8.5, 30 % PEG 300. For cryoprotection, a drop of reservoir solution was added to the crystals, which were then maintained in liquid nitrogen.

### Diffraction data collection and structure determination

Datasets were collected using synchrotron radiation (λ = 1.0 Å) at the X06SA-beamline (Swiss Light Source, Villingen, Switzerland). X-ray intensities were assessed and data reduction carried out with XDS and [32]. A dataset for pSer111-Rab8A:GDP was collected from a single crystal, to a resolution of 2.4 Å. For pSer111-Rab8A:GppNHp, two partial datasets from two crystals were merged to produce a dataset at 2.5 Å resolution (Table 1). Initial phases were calculated for pSer111-Rab8A:GDP and pSer111-Rab8A:GppNHp by Patterson search procedure using the deposited coordinates of Rab8•GDP (PDB ID: 4LHV) and Rab8•GppNHp (PDB ID: 4LHW), respectively [21]. Molecular replacement was carried out using PHASER [33]. Iterative model rebuilding and refinement were performed with COOT [34] and phenix.refine [35], applying non-crystallographic symmetry restraints and Translation/Liberation/Screw tensors. Eventually, water molecules were placed automatically using ARP/wARP [36]. The final models of pSer111-Rab8A:GDP (PDB ID: 6STF) and pSer111-Rab8A:GppNHp (PDB ID: 6STG) converged to R_work_/R_free_ values of 23.3/26.2 and 24.3/27.8. Coordinate geometries were confirmed via MOLPROBITY [37] to possess good stereochemistry and small bond-length and angle RMSDs, with no residues lying in the disallowed region of the Ramachandran plot (Table 1).

**Table 1:** Crystallographic data collection and refinement statistics.

|  | p-Ser111-Rab8A:GDP | p-Ser111-Rab8A:GppNHp |
| --- | --- | --- |
| **<u>Crystal parameters</u>** |  |  |
| Space group | C2 | P2<sub>1</sub> |
| Cell constants | a = 117.6 Å; b = 75.2 Å;<br>c = 107.7 Å; β = 100.5 Å | a = 34.9 Å; b = 93.7 Å;<br>c = 56.1 Å; β = 103.0 Å |
| Subunits / AU<sup>a</sup> | 5 | 2 |
| **<u>Data collection</u>** |  |  |
| Beam line | X06SA, SLS | X06SA, SLS |
| Wavelength (Å) | 1.0 | 1.0 |
| Resolution range (Å)<sup>b</sup> | 50 – 2.4 (2.5 – 2.4) | 50 – 2.5 (2.6 – 2.5) |
| No. observed reflections | 105949 | 35644 |
| No. unique reflections<sup>c</sup> | 35267 | 11734 |
| Completeness (%)<sup>b</sup> | 97.2 (96.9) | 95.9 (92.3) |
| R<sub>merge</sub> (%)<sup>b, d</sup> | 0.096 (0.574) | 0.123 (0.578) |
| I/σ (I)<sup>b</sup> | 9.5 (2.1) | 6.5 (2.0) |
| **<u>Refinement (REFMAC5)</u>** |  |  |
| Resolution range (Å) | 30 – 2.4 | 30 – 2.5 |
| No. refl. working set | 33488 | 11143 |
| No. refl. test set | 1763 | 586 |
| No. non hydrogen | 6875 | 2845 |
| No. of pSer111 | 5 | 2 |
| No. of Mg<sup>2+</sup> | 5 | 2 |
| No. of nucleotide | 5 | 2 |
| Solvent | 45 | 17 |
| R<sub>work</sub> / R<sub>free</sub> (%)<sup>e</sup> | 23.3 / 26.2 | 24.3 / 27.8 |
| r.m.s.d. bond (Å) / (°)<sup>f</sup> | 0.002 / 1.2 | 0.002 / 1.2 |
| Average B-factor (Å<sup>2</sup>) | 48.4 | 51.8 |
| Ramachandran Plot (%)<sup>g</sup> | 97.8 / 2.2 / 0 | 98.5 / 1.5 / 0 |
| PDB accession code | 6STF | 6STG |

<sup>[a]</sup> Asymmetric unit
<sup>[b]</sup> The values in parentheses for resolution range, completeness, R<sub>merge</sub> and I/σ (I) correspond to the highest resolution shell
<sup>[c]</sup> Data reduction was carried out with XDS. Friedel pairs were treated as identical reflections
<sup>[d]</sup> $R_{\text{merge}}(I) = \sum_{hkl} \sum_j |I(hkl)_j - \langle I(hkl) \rangle| / \sum_{hkl} \sum_j I(hkl)_j$ , where $I(hkl)_j$ is the $j^{\text{th}}$ measurement of the intensity of reflection hkl and $\langle I(hkl) \rangle$ is the average intensity
<sup>[e]</sup> $R = \sum_{hkl} | |F_{\text{obs}}| - |F_{\text{calc}}| | / \sum_{hkl} |F_{\text{obs}}|$ , where R<sub>free</sub> is calculated for a randomly chosen 5% of reflections, which were not used for structure refinement, and R<sub>work</sub> is calculated for the remaining reflections
<sup>[f]</sup> Deviations from ideal bond lengths / angles
<sup>[g]</sup> Percentage of residues in favored / allowed / outlier region

### NMR experiments and analysis

Uniformly ^15^N labelled Rab1B was expressed and purified as mentioned in the method section, with M9 minimal medium containing ^15^NH_4_Cl (1 g/l) as sole nitrogen source. For phosphorylated Rab1B, unlabeled phosphorylated-serine was added to the media since the isotopically labelled derivative is not available. NMR samples were exchanged into final NMR buffer consisting of 20 mM sodium phosphate, 50 mM NaCl, 1mM DTT, 1mM MgCl_2_, pH 6.5 with respective 10µM GDP or GppNH. 10% ^2^H_2_O was added for deuterium lock. NMR sample concentrations were 100 to 300µM range. Two dimensional-proton ^1^H, ^15^N NMR water-flipback WATERGATE HSQC experiments were recorded on a 600MHz Bruker NMR spectrometer equipped with a TCI cryoprobe. Experiments were recorded with acquisition times of 122msec and 62msec in the ^1^H and ^15^N dimensions, with a interscan relaxation delay of 1 second. All measurements were performed at 298K. Proton chemical shifts were calibrated with respect to DSS (2,2-dimethyl-2-silapentane-5-sulfonate), while nitrogen chemical shifts were calibrated indirectly. For data processing, all FID were multiplied with a square-shifted-sine-bell window function and data points with zero-filling before Fourier transformation using the Bruker TOPSPIN3.5pl7 software. Spectra were analysed using CCPN v.2.4.2 [38].

### Cell Culture

Flp-In TREx HEK293 cells expressing PINK1-3xFLAG were cultured using DMEM (Dulbecco’s modified Eagle’s medium) supplemented with 10 % FBS (foetal bovine serum), 2 mM L-Glutamine, 1X Pen/Strep and 1X non essential amino acids (Life Technologies). Flp-In T-Rex HeLa stable cell lines were cultured in the above mentioned media supplemented with 15 μg/ml of Blasticidin and 100 μg/ml of Hygromycin. Cell transfections of untagged LRRK2 [R1441G], GFP-Rab8A WT or S111A were performed using polyethylene method [39]. Cultures were induced to express protein by addition of 0.1 μg/ml of Doxycycline to the medium for 24 h. To uncouple mitochondria, cells were treated with 10 μM CCCP (Sigma) dissolved in DMSO for 3 hours.

### Immunoblotting

Samples were subjected to SDS-PAGE and transferred onto nitrocellulose membranes. Membranes were blocked with 5 % BSA in TBS-T and incubated at 4 °C overnight with the indicated antibodies, diluted in 5 % BSA. Membranes for kinase assay screen were incubated with HRP-conjugated secondary antibodies (1:10,000) in 5 % milk for 1 h at room temperature. Membranes were exposed with ECL substrate. Additional kinase assay immunoblotting membranes were incubated with secondary antibodies, conjugated with LICOR IRDye in TBS-T (1:10,000) and imaged using the LICOR Odyssey software.

### Antibodies

The following antibodies were produced by the MRCPPU Reagents and Services at the University of Dundee in sheep: anti-Rab8A phospho-Ser111 (S503D), 4^th^ bleed; raised against residues 104-117 of human Rab8A: RNIEEHApSADVEKMR), Anti Rab8A pThr72 (S874D), raised against residues 65-79 of human Rab8A: AGQERFRpTITTAYYR, Anti Rab1B pSer111 (S817D), 3^rd^ bleed; raised against residues 104-118 of human Rab1B: CQEIDRYApSENVNKLR. Additional phospho-Rab antibodies were produced in development with Abcam: Anti-Rab8A phospho-Thr72 (ab230260 MJFF-20). Anti-Rab8A (D22D8 #6975) was from Cell Signalling Technologies, Anti-6X His Tag antibody (ab18184) from Abcam and Anti-LRRK2 C-terminal (Dardarin clone N241A/34) from NeuroMab. Anti-Rab1B phospho-Thr72 (ABS2131) was from Merck Millipore. Antibodies were used at a final concentration of 1 µg/ml, diluted in 5 % BSA.

## Results

### Identification of MST3 and TAK1 as direct kinases of Rab8A Thr72

Previous biochemical analysis undertaken by our laboratory has demonstrated that PINK1 does not directly phosphorylate Rab GTPases at Ser111 [8]. To search for candidate Rab Ser111-directed kinases, we undertook an *in vitro* [γ-^32^P]-ATP-based screen of ∼140 recombinant kinases against recombinant wild-type (WT) or Ser111Ala (S111A) Rab8A expressed in the GDP-bound conformation. We identified multiple kinases capable of phosphorylating wild-type Rab8A *in vitro* as judged by autoradiography including LRRK2 that has previously been reported to phosphorylate Rab8A at Thr72 (Figure 1A) [10]. Immunoblotting of reactants with a phospho-Ser111-Rab8A (pSer111-Rab8A) antibody did not reveal any kinases that directly phosphorylated GDP-bound Rab8A at Ser111 (Supplementary Figure 1) and consistent with this, we did not identify any kinases that differentially phosphorylated the WT Rab8A more efficiently than the S111A Rab8A mutant (Supplementary Figure 1). Previous studies have suggested that the targeting of kinases by Rabs can be influenced by their nucleotide status e.g. LRRK2 preferentially phosphorylates the GDP-bound form of Rab8A [10]. We therefore repeated the *in vitro* screen with GTP-bound Rab8A, however, under these conditions, we did not identify any kinase directed towards Ser111 by immunoblotting with a pSer111-Rab8A antibody (Supplementary Figure 2).

**Figure 1.**
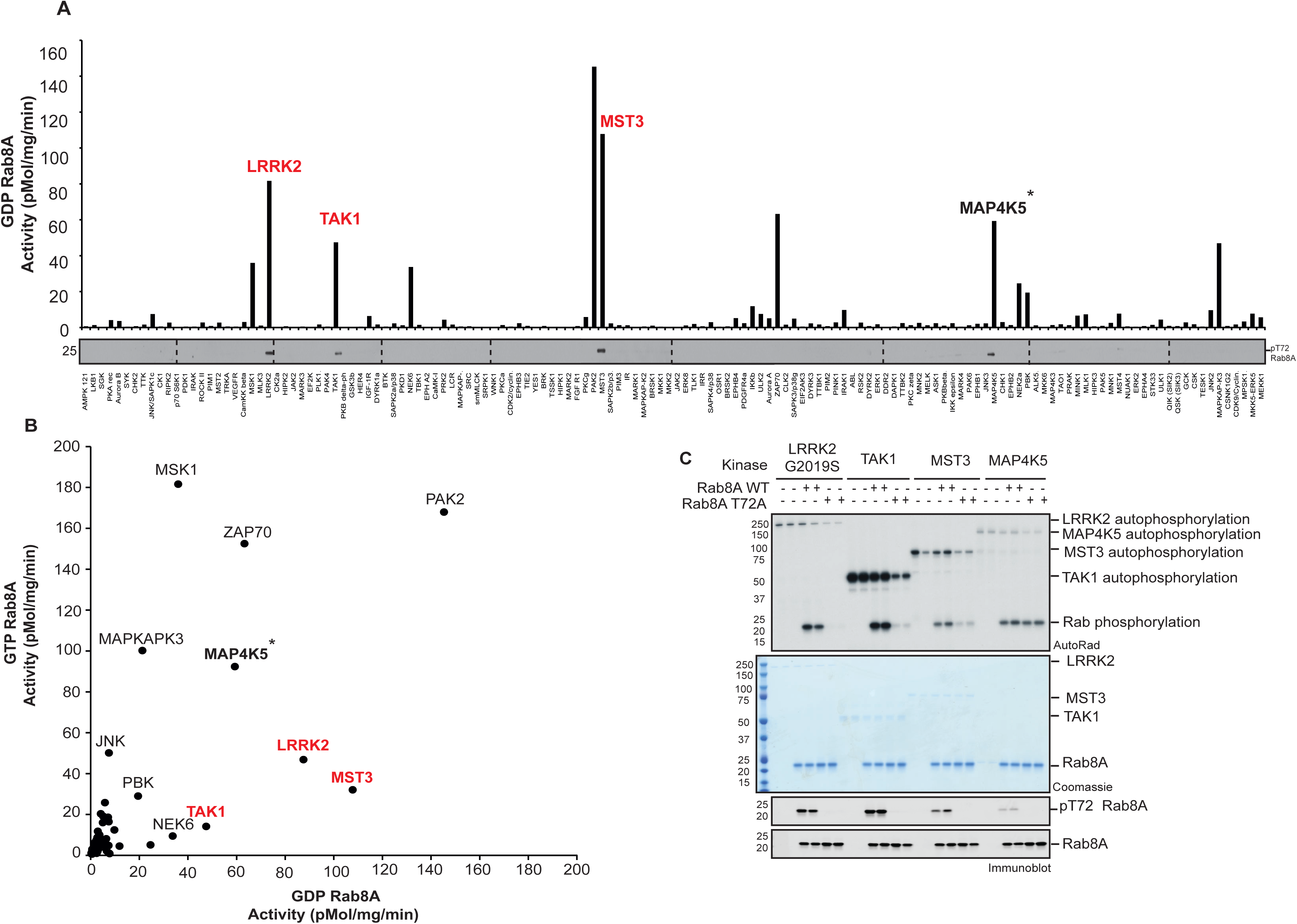
TAK1 and MST3 directly phosphorylate Rab8A at Thr72. **A:** Summary analysis of kinase assay screen quantification to identify novel Rab8A kinases. 140 protein kinases from the MRC PPU Reagents and Services library were assessed for the phosphorylation of GDP-Rab8A. 100 ng of protein kinase was incubated with 2 µg substrate and [γ-^32^P] ATP for 30 min. Samples were subjected to SDS-PAGE and analysis with Cerenkov counting of Coomassie substrate bands, displayed graphically. Full kinase assay analysis by autoradiography and immunoblotting is described in Supplementary Figure 1. Proteins highlighted in red were identified to phosphorylate Rab8A at Thr72 by phospho-specific immunoblotting. **\***Thr72 is minor MAP4K5 phosphorylation site and occurs in GDP-Rab8A but not GTP-Rab8A conformation. **B:** Correlative analysis of kinase phosphorylation of GTP-Rab8A (y-axis) versus GDP-Rab8A (x-axis). The GTP-Rab8A kinase screen was performed in a similar manner to GDP-Rab8A kinase screen, with full analysis described in Supplementary Figure 2. Kinases that phosphorylate Thr72 as the major site of Rab8A are highlighted in red. **C.** Mutation of Thr72Ala (T72A) reduces Rab8A phosphorylation by LRRK2, TAK1 and MST3 but not MAP4K5. The indicated kinases were incubated in the presence of WT or T72A GDP-Rab8A. Samples were subjected to SDS-PAGE with analysis by Coomassie staining and [γ-^32^P] incorporation as visualised by autoradiography and immunoblotting analysis using the indicated antibodies.

Whilst we did not identify the Ser111 kinase in these screens, immunoblotting of reactants with a phospho-Thr72-Rab8A (pThr72-Rab8A) antibody revealed that three kinases, TAK1, MST3 and MAP4K5 were capable of phosphorylating Rab8A at Thr72 equivalent to the LRRK2 site (Figure 1A and Supplementary Figure S1 & S2). Similar to LRRK2, there appeared to be a preference for TAK1 and MST3 to phosphorylate GDP-bound Rab8A compared to GTP-bound Rab8A (Figure 1B and Supplementary Figure 1 & 2). Whilst MAP4K5 phosphorylated Thr72 in GDP-bound Rab8A but not GTP-bound Rab8A, the overall phosphorylation of Rab8A was unchanged indicating the presence of additional non-Thr72 phosphorylation sites targeted by MAP4K5 *in vitro* (Figure 1A-B and Supplementary Figure 1 & 2). Mutating Thr72 to Ala substantially reduced the phosphorylation of Rab8A by TAK1 and MST3 thereby confirming that this residue represents the major site of phosphorylation (Figure 1C). In contrast there was no reduction in phosphorylation by MAP4K5 indicating that the Thr72 residue is likely to be a minor phosphorylation site (Figure 1C). The screen also revealed other kinases including PAK2 and ZAP70 that can efficiently phosphorylate Rab8A (Figure 1A-B and Supplementary Figure 1 & 2) and in future work it would be interesting to determine the site(s) of phosphorylation and understand how these affect Rab8A function.

### Phosphorylation of Rabs at Ser111 impairs interactions with Switch II binding regulatory proteins

Upon attachment to their target membranes, Rab GTPases act as molecular switches that are activated by a guanine nucleotide exchange factor (GEF) that catalyses the exchange of the bound guanosine-5’-diphosphate (GDP) for the more abundant guanosine-5’-triphosphate (GTP). Once active the GTP-bound Rab can interact with specific effector proteins which mediate a variety of downstream functions including binding to cargoes, enabling vesicle formation, and promoting vesicle fusion to target membranes [12]. Rabs are inactivated by GTPase activating proteins (GAPs) that promote the hydrolysis of GTP to GDP and the inactive Rab is then relocalised to the cytosol [12]. Structural analysis of Rab8A reveals that the PINK1-regulated Ser111 site lies within a short stretch of amino acids within the C-terminal region termed the RabSF3 motif [13]. The RabSF3 motif of Rab8A lies close to the switch II region that mediates GEF binding and we have previously observed that a Ser111Glu phosphomimetic (S111E) mutant of Rab8A negatively affects its ability to be activated by its cognate GEF, Rabin8 [8]. However, phosphomimetic substitutions often fail to recapitulate the full functional effect of phosphorylation and have also been reported to render Rabs non-functional [14].

We therefore investigated the effect of the phosphorylation at Ser111 on Rab8A using preparative phosphorylated Rab8A. Employing previously published genetic code expansion technologies for the incorporation of pSer65 into Ubiquitin [15], we were able to produce pSer111-Rab8A in the inactive GDP-bound conformation (pSer111-Rab8A:GDP) and the active GTP-bound conformation (pSer111-Rab8A:GppNHp) with a yield of 0.15 mg per liter of bacterial culture (Supplementary Figure 3). The purity of the proteins was confirmed by SDS-PAGE (Supplementary Figure 3A). Thermal shift analysis demonstrated no significant difference in melting points between pSer111-Rab8A:GDP and WT-Rab8A:GDP or between pSer111-Rab8A:GppNHp and WT-Rab8A:GppNHp suggesting that the pSer111-Rab8A exhibited similar protein stability to the WT-Rab8A (Supplementary Figure 3C). To assess the stoichiometry of phosphorylation, we initially employed Phos-tag™ acrylamide SDS-PAGE to probe for the presence of pSer111-Rab8A:GDP and pSer111-Rab8A:GppNHp relative to the non-phosphorylated (WT) form. Both proteins, pSer111-Rab8A:GDP and pSer111-Rab8A:GppNHp, completely shifted +10kDa compared to WT-Rab8A:GDP and WT-Rab8A:GppNHp indicating stoichiometric phosphorylation (Supplementary Figure 3B). LC-MS analysis demonstrated that the mass spectra of pSer111-Rab8A showed the expected increase in molecular weight of +80 Da confirming the presence of a single phosphorylation (Supplementary Figure 3D). There was no trace of the non-phosphorylated (WT) form in the mass spectra of pSer111-Rab8A suggesting that the Rab8A protein produced by genetic code expansion was quantitatively phosphorylated (Supplementary Figure 3D).

We next assessed the interaction of phosphorylated Rab8A with its GEF and GAP by measuring the nucleotide exchange and GTP hydrolysis rate respectively. We first compared the Rabin8-mediated GDP-GTP exchange and the TBC1D20-mediated GTP hydrolysis rate in WT-Rab8A and pSer111-Rab8A measuring the change in their intrinsic tryptophan (Trp) fluorescence [16] upon nucleotide exchange over time (Figure 2A-B). We observed an 80% reduction of the GEF activity (Figure 2A and B) and the GAP activity was reduced by approximately 60% (Figure 2A and B).

**Figure 2:**
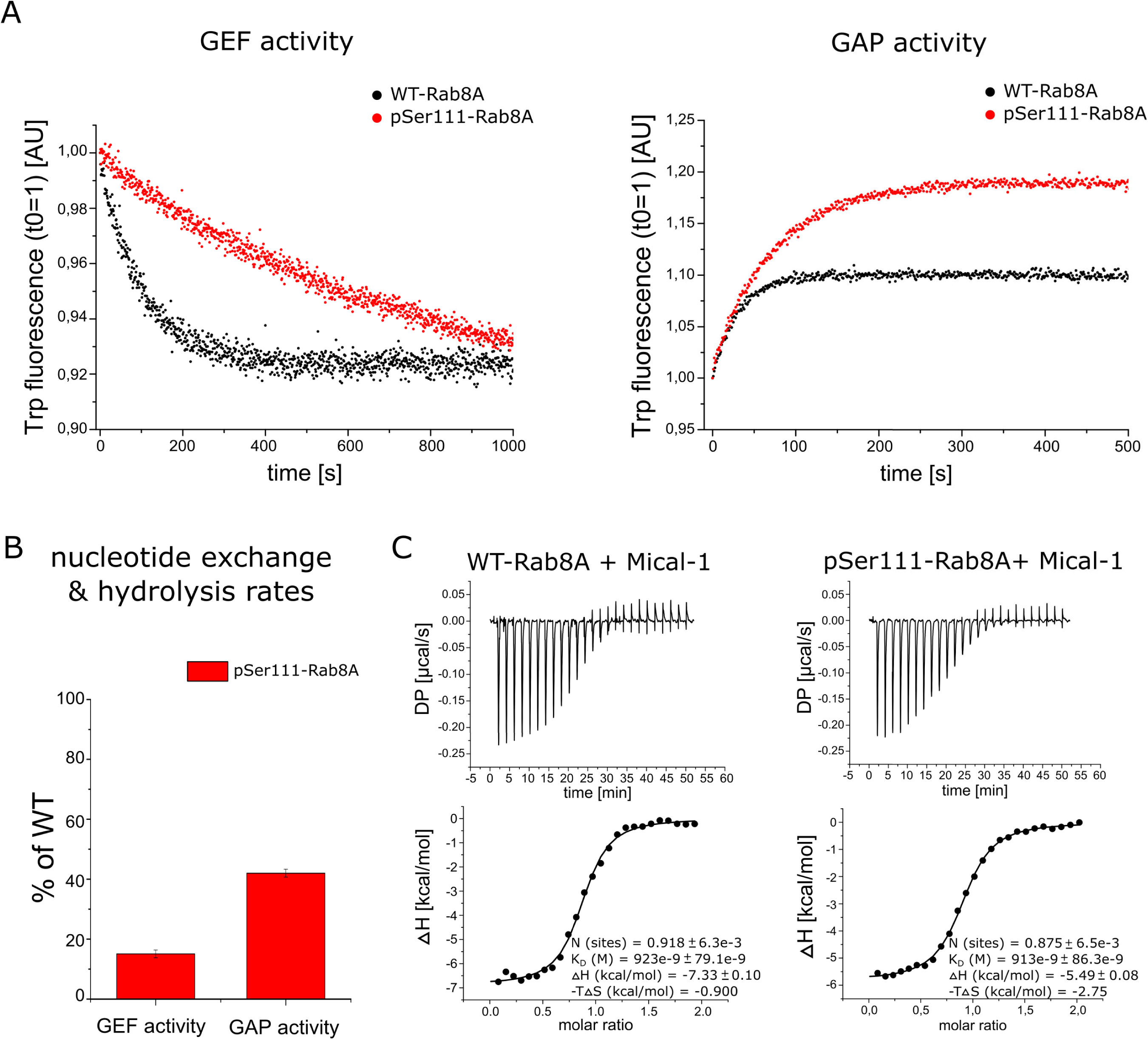
Rab8A Ser111 phosphorylation impairs GEF and GAP interactions. **A.** (Right Panel) The GDP-GTP exchange catalyzed by the GEF Rabin8 (GEF activity), and the (Left Panel) GTP hydrolysis catalyzed by the GAP TBC1D20 (GAP activity) were monitored for WT-Rab8A-His_6_ (black) and pSer111-Rab8A-His_6_ (red) proteins via intrinsic tryptophan (Trp) fluorescence. Data are representative for three independent experiments. **B.** The nucleotide exchange (GEF activity) and GTP hydrolysis rates (GAP activity) were determined using a single exponential fit, and are represented as % of WT. All data (n=3) are represented as mean ± S.D. **C.** The binding of the effector Mical-1 to WT-Rab8A-His6 and pSer111-Rab8A-His6 was determined by isothermal titration calorimetry (ITC). Data are representative for two independent experiments. The dissociation constants (K_D_) for binding affinity are indicated in each graph.

Rab1B shares many structural similarities with Rab8A [13] and can also be phosphorylated in a PINK1-dependent manner in cells at Ser111 equivalent to the Rab8A site (Supplementary Figure 4) [8]. We therefore also expressed and purified preparative wild-type (WT-Rab1B:GDP) and Ser111-phosphorylated GDP-bound Rab1B (pSer111-Rab1B:GDP) and wild-type (WT-Rab1B:GDP) and Ser111-phosphorylated GTP-bound Rab1B (pSer111-Rab8A:GppNHp) (Supplementary Figure 3A-D). A similar trend was observed for pSer111-Rab1B, whereby the GEF activity was decreased by 60% and the GAP activity by 40% (Supplementary Figure 5).

To determine whether Ser111 phosphorylation was mediating specific effects via the Switch II domain we determined the binding affinity of WT- and pSer111-Rab8A GppNHp-bound forms with the effector Mical-1 which does not directly bind via the Switch II domain [17] (Supplementary Figure 6). Using isothermal titration calorimetry (ITC), we obtained a K_D_ value of 923 nM for the binding affinity of WT-Rab8A and Mical-1 and a K_D_ of 913 nM for pSer111 Rab8A with Mical-1 suggesting that the phosphorylation at Ser111 has no effect on Rab8-Mical-1 effector interaction (Figure 2C). The K_D_ values obtained from the ITC measurements are more than 15 times higher compared to a previous study from Rai and colleagues [18] characterizing the interaction between WT-Rab8A and Mical-1, which may be due to the C-terminal His_6_ tag on our WT-Rab8A proteins lowering the binding affinity towards Mical-1. Overall our data suggests that the phosphorylation of Rab8A at Ser111 may influence Switch II binding by regulators and disrupts interactions with its cognate GEF and moderately impairs its interaction with GAPs.

### Phosphorylation of Rabs at Ser111 disrupts the ability of LRRK2 to phosphorylate Rabs at Thr72

The LRRK2 kinase has previously been shown to phosphorylate Rab8A at position Thr72 within the switch II domain [10], however, the structural determinants and mechanism by which LRRK2 phosphorylates Rabs remains unknown. Since Rab8A Ser111 phosphorylation can disrupt switch II interactions mediated by Rabin8 and TBC1D20, we next investigated whether Ser111 phosphorylation may impact on LRRK2 mediated phosphorylation of Rab8A at Thr72. To address this we performed an *in vitro* kinase assay using recombinant WT GDP or GTP-bound Rab8A (WT-Rab8A:GDP; WT-Rab8A:GppNHp) or pSer111 GDP or GTP-bound Rab8A (pSer111-Rab8A:GDP; pSer111-Rab8A:GppNHp) incubated with [γ-^32^P]-ATP and an activating LRRK2 mutant [G2019S]. Consistent with previous studies we observed that LRRK2 [G2019S] efficiently phosphorylated WT-Rab8A:GDP but was less able to phosphorylate WT-Rab8A:GppNHp as determined by measuring [γ-^32^P]-ATP incorporation via autoradiography and by immunoblotting of reactants with a pThr72 Rab8A phospho-specific antibody (Figure 3A) [10]. Strikingly we observed that the ability of LRRK2 [G2019S] to phosphorylate pSer111-Rab8A:GDP was disrupted compared to WT-Rab8A:GDP and a similar reduction was also observed for pSer111-Rab8A:GppNHp compared to WT-Rab8A:GppNHp (Figure 3A). This was not due to inhibition of LRRK2 catalytic activity since LRRK2 autophosphorylation was unchanged (Figure 3A). It has previously been shown that LRRK2 can phosphorylate Rab1B *in vitro* at Thr72 equivalent to the Rab8A Thr72 site [10]. To determine whether the effect of Ser111 phosphorylation was more generalizable to other LRRK2 Rab substrates that are common to PINK1, we next assessed the ability of LRRK2 [G2019S] to phosphorylate Rab1B *in vitro*. Similar to Rab8A, we observed a significant reduction in the ability of LRRK2 [G2019S] to phosphorylate pSer111-Rab1B:GDP or pSer111-Rab1B: GppNHp compared to WT-Rab1B as judged by autoradiography and immunoblotting of reactants with a p-T72 Rab1B phospho-specific antibody (Supplementary Figure 7A).

**Figure 3:**
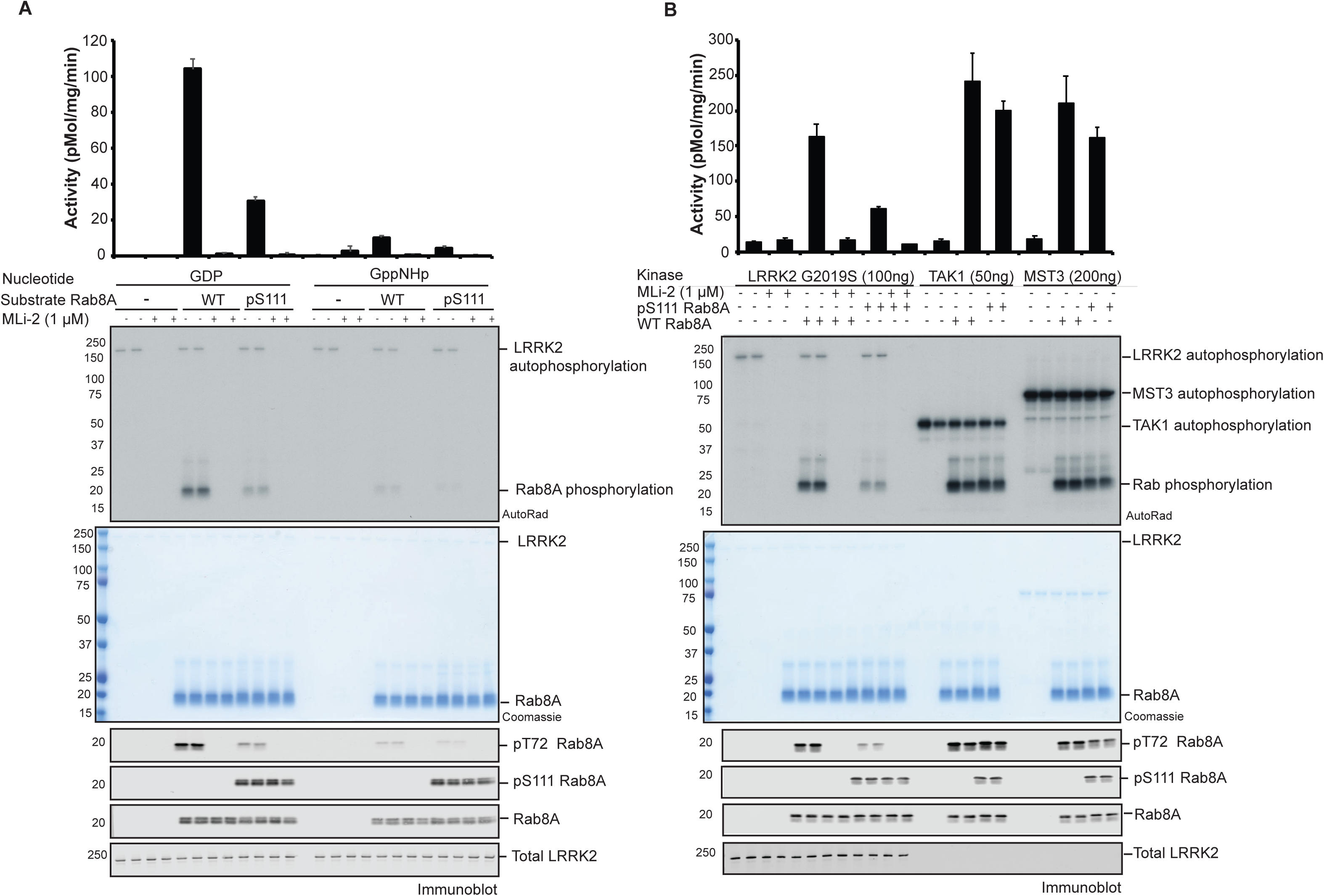
Rab8A Ser111 phosphorylation disrupts LRRK2-mediated Thr72 phosphorylation *in vitro*. **A.** 100 ng of recombinant LRRK2 [G2019S] protein was incubated with [γ-^32^P] and 2 µg of WT or pSer111 Rab8A, in either the GDP or GTP (GppNHp) bound conformation, in the absence or presence of the LRRK2 inhibitor, MLi-2 for 30 min. Samples were subjected to SDS-PAGE, and analysed by either Coomassie staining, [γ-^32^P] incorporation measured by autoradiography with Cerenkov counting (top panel) and immunoblot analysis, using the indicated antibodies (lower panel). Results are means +/- S.E.M. (*n*=4). **B.** Thr72 specific kinases were assessed for phosphorylation of GDP bound WT or pSer111 Rab8A. 100 ng LRRK2 G2019S, 50 ng TAK1 or 200 ng MST3 were incubated with 2 µg substrate using identical conditions and analysis as in panel A.

We next compared the ability of other Rab Thr72-directed kinases, MST3 and TAK1, to phosphorylate pSer111 Rab8A. We performed a side-by-side comparison of the efficiency of phosphorylation of WT-Rab8A:GDP or pSer111-Rab8A:GDP by LRRK2, MST3 and TAK1 (Figure 3B). As previously observed, we found that pSer111-Rab8A:GDP reduced LRRK2-mediated phosphorylation compared to WT-Rab8A:GDP. In contrast, MST3 and TAK1 were still capable of phosphorylating pSer111-Rab8A:GDP to a similar extent as WT-Rab8A:GDP (Figure 3B). Consistent with these results we also observed no effect of pSer111-Rab1B:GDP on the ability of MST3 and TAK1 kinases to phosphorylate Thr72 compared to WT-Rab1B:GDP compared to LRRK2 (Supplementary Figure 7B).

### Phosphorylation at Ser111 does not induce major structural changes in Rab8A and Rab1B

To understand the mechanism by which Ser111 phosphorylation impacts on its interactions with its GEF, GAP and LRRK2, we next sought to solve the structure of pSer111-Rab8A. We were able to obtain well-diffracting crystals for both pSer111-Rab8A:GDP (2.4 Å resolution, R_free_ = 26.2%, PDB ID: 6STF) and pSer111-Rab8A:GppNHp (2.4 Å resolution, R_free_ = 27.8%, PDB ID: 6STG) (Table 1). Phosphorylated Rab8A exhibits a typical small GTPase fold [19, 20] with highly flexible Switch regions in the GDP-bound form, and well-defined structured Switch regions in the GppNHp-bound form (Figure 4). The pSer111 residue is located C-terminal of the helix α3 in a loop above the Switch II region (α2). The superimposed structures of pSer111-Rab8A and WT-Rab8A [21] showed that the phosphorylation at Ser111 does not impact the overall structure of Rab8A (Figure 4 and Supplementary Figure 8). Nevertheless, the spatial proximity of pSer111 to the Switch II region (α2) could allow for dynamic interactions between pSer111 and residues within Switch II that cannot be detected by crystallography. In order to test the potential conformational consequences of Ser111 phosphorylation, we performed nuclear magnetic resonance (NMR) experiments and compared the phosphorylated and non-phosphorylated forms. However, since we were unable to produce isotopically labelled phosphorylated Rab8A, we instead produced the close homologue Rab1B as a surrogate (Supplementary Figure 9). When comparing the 1H 15N Heteronuclear Single Quantum Coherence (HSQC) NMR data for Rab1B pSer111 in the GDP and GppNHp states, most of the chemical shifts overlap very well, indicating that Ser111 phosphorylation, probably does not lead to major structural alterations on Rab1B in solution. However, some of the resonances show shifts because of a change in the chemical environment close to the phospho group (Supplementary Figure 10). Overall, the lack of conformational changes observed in the crystal structures (pSer111-Rab8A) are consistent with the lack of structural changes seen in solution (pSer111-Rab1B). In future work it would be interesting to employ NMR to solve the structure of Rab1B in complex with its effectors to determine how pSer111 might alter the interaction between the Switch II region and known Rab1B regulators such as the GEF, as well as LRRK2.

**Figure 4:**
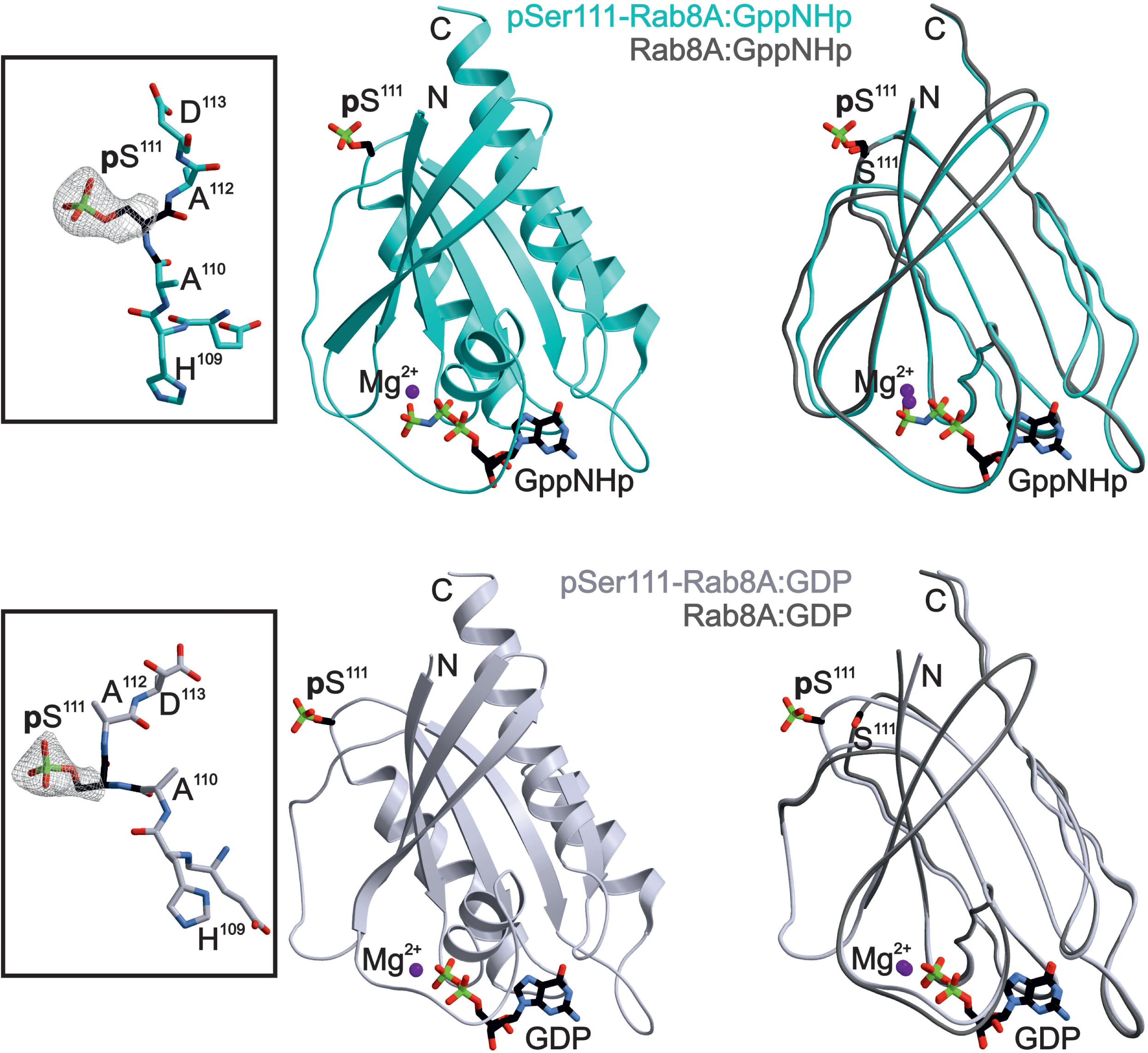
X-ray structure of Rab8A phosphorylated at Ser111. **A.** Ribbon drawings of active pSer111-Rab8A:GppNHp (PDB ID 6STG, cyan, upper panel) and inactive pSer111-Rab8A:GDP proteins (PDB ID 6STF, lavender, lower panel), respectively. pSer111 and nucleotides are shown as balls and sticks model; Mg^2+^ is illustrated as a purple sphere. The inset on the left depicts the 2F_o_-F_c_ electron density map of the pSer111 side chain. The panel on the right displays the overlay of the phosphorylated Ser111-Rab8A structure with its unmodified counterpart (Rab8A:GppNHp, PDB ID 4LHW; Rab8A:GDP, PDB ID 4LHV) [21].

### Evidence of interplay of PINK1 and LRRK2 signalling on Rab8A in cells

Under basal conditions in which PINK1 is inactive, Rab8A is phosphorylated by LRRK2. Previous studies have demonstrated that the hyperactive mutant LRRK2 [R1441G] leads to increased Rab8A Thr72 phosphorylation [9, 10]. Under mitochondrial depolarisation that can be induced by mitochondrial uncouplers, (e.g. CCCP) PINK1 is stabilized and activated and can phosphorylate Rab8A at Ser111 [8]. We therefore expressed LRRK2 [R1441G], and wild-type or a non-phosphorylatable S111A mutant of GFP-Rab8A in HEK 293 Flp-In TREx cells stably expressing PINK1-3xFLAG. Cells were pre-treated with DMSO or 100 nM MLi-2 (LRRK2 inhibitor) for 1.5 h followed by either 10 μM CCCP or DMSO control for 3 h. Cells were lysed and whole cell extracts analysed by immunoblotting with anti-GFP, and anti-LRRK2 antibodies that confirmed uniform expression across all conditions (Supplementary Figure 11). Immunoblotting with anti-PINK1 antibodies and OPA1 antibodies confirmed equivalent expression across basal and mitochondrial depolarized conditions (Supplementary Figure 11). To visualize phosphorylated Rab8A we undertook Phos-tag™ SDS-PAGE and immunoblotting with anti-GFP antibody, demonstrating a single electrophoretic band shift of ∼40% total Rab8A under basal conditions that was abolished by MLi-2 treatment indicating LRRK2-mediated Thr72 phosphorylation (Figure 5 and Supplementary Figure 11). Furthermore, this was confirmed by an anti-phospho-Rab8A-Thr72 antibody (Figure 5 and Supplementary Figure 11). Upon combined treatment of MLi-2 and CCCP, we observed the appearance of a single electrophoretic band of ∼20% Rab8A due to PINK1-mediated Ser111 phosphorylation that migrated at a similar pattern to LRRK2-mediated monophosphorylated Rab8A and was also positively detected by the anti-phospho-Rab8A-Ser111 antibody (Figure 5 and Supplementary Figure 11). Upon removal of MLi-2 and in the presence of CCCP, we observed a double electrophoretic band shift with the upper band representing the dual phosphorylated Rab8A as confirmed by immunoblotting of the Phos-tag™ gel with anti-phospho-Rab8A-Thr72 and anti-phospho-Rab8A-Ser111 antibodies (Figure 5 and Supplementary Figure 11). Immunoblotting with anti-pSer111 antibody revealed equivalent levels pSer111 in the monophosphorylated Rab8A or double phosphorylated Rab8A species suggesting that there is no preference for the incorporation of pSer111 into unmodified or pThr72-Rab8A (Figure 5). Thus, the presence of the Thr72 phosphorylation does not appear to inhibit the phosphorylation of Rab8A at position Ser111 by PINK1. However, overall, the double phosphorylated species represents only ∼2% of the total Rab which is less than predicted based on Ser111-monophosphorylation levels and suggests that pThr72 is preferentially incorporated into unmodified Rab8A rather than Ser111-phosphorylated Rab8A consistent with our *in vitro* studies (Figure 5).

**Figure 5.**
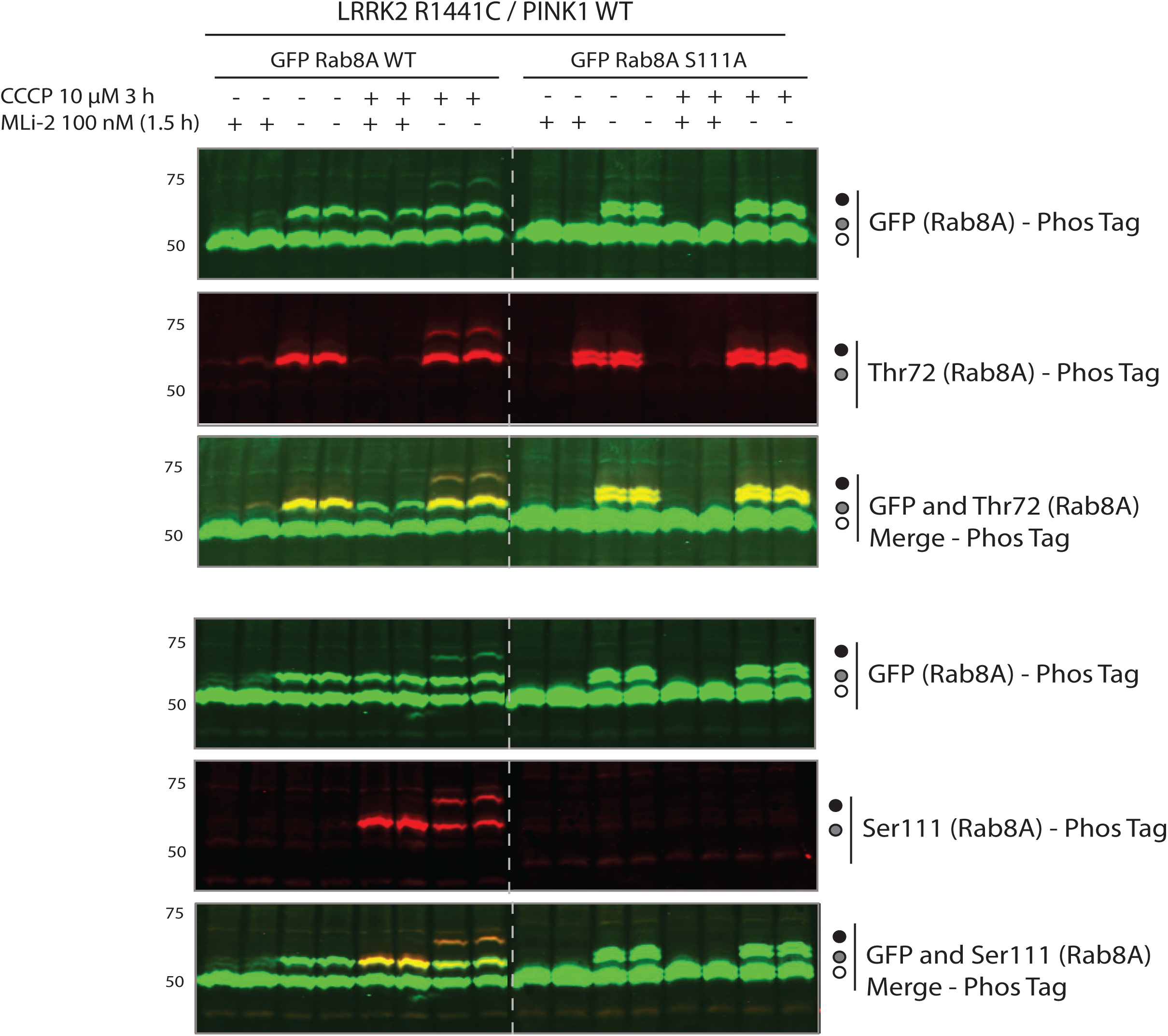
PINK1 and LRRK2 signalling converge on Rab8A in cells. HEK293 Flp-In TREx cells stably expressing PINK1-3XFLAG were co-transfected with 9 μg of LRRK2 R1441G and 3 μg of GFP Rab8A WT or S111A. Cells were treated with DMSO, 100 nM MLi-2 (1.5 h), 10 μM CCCP (3 h) or a combination of each. Cells were lysed and underwent Phos-tag analysis with total and phospho-specific antibodies using the LICOR Odyssey imaging system for detection. IRDye 800CW fluorescent secondary antibody was utilised for GFP Rab total signal, whilst IRDye 680RW fluorescent antibody was used for the specific phospho-Rab detection. Presence of yellow fluorescence upon multiplexing with Image Studio software, indicates the presence of site-specific phosphorylated Rab within the GFP total Rab population.

## Discussion

Using genetic code expansion technology, we have been able to demonstrate that PINK1-dependent phosphorylation of Rab GTPases at Ser111 impairs the ability of Rabs to interact with GEFs and GAPs. Interference of the GEF or GAP binding would trap the phosphorylated Rab protein in either the inactive GDP- or active GTP-bound state (Figure 2 and Supplementary Figure 5). Furthermore, we have also found that PINK1-dependent Rab phosphorylation blocks the ability of the LRRK2 kinase but not TAK1 or MST3 to phosphorylate Thr72 (Figures 3, 6 and Supplementary Figure 7).

The Ser111 residue lies within a C-terminal *α*3/*β*5 loop of Rab8A, that connects the *α* helix 3 with the *β* sheet 5 known as the SF3 motif and lies adjacent to the Switch II domain (Figure 4) [22, 23]. The Switch II region represents the major interaction surface for Rab effectors and undergoes dynamic conformational change upon alteration between its GTP and GDP-bound forms [13]. Across all Rab structures in their active state, there is structural heterogeneity both within the switch domains, and also the SF3 motif (with little change in other regions) and together these regions explain how different Rab proteins can bind to specific sets of effector proteins to regulate distinct pathways [24–26]. Previous studies have reported a role for the SF3 motif in selective recruitment of effectors via direct effector-specific contact sites e.g. the recruitment of Rabphillin by Rab3a via SF3 motif binding sites [22]. However, the role of post translational modifications including phosphorylation within the SF3 motif on effector interactions has not been studied with preparative amounts of modified Rab proteins.

The phosphorylation at Ser111 did not induce any major changes in the structure of Rab8A (Figure 4 and Supplementary Figure 8), nor in Rab1B (Supplementary Figure 9 and 10), despite impacting on GEF/GAP interactions (Figure 2A-B and Supplementary Figure 5). The effect of pSer111 on Rabin8-mediated nucleotide exchange is similar in extent to the previously analyzed phosphomimetic S111E-Rab8A mutant, suggesting that the introduction of negative charges within the SF3 motif may influence GEF/GAP binding [8]. Consistent with this possibility, the Rab8A:Rabin8 structure has recently been analyzed using Molecular Dynamics (MD) simulation revealing a salt-bridge interaction between residue Asp187 of Rabin8 with Arg79 that lies within the Switch II domain of Rab8A [27]. Ser111 is located opposite a negative surface patch of Rabin8 in which Asp187 lies and addition of a negative charge at Ser111 either by S111E or pSer111 weakens/disrupts the interaction between Asp187 of Rabin8 and Arg79 of the Rab8A [27]. Whilst the Arg79 residue is not part of the TBC1D20 binding site in Rab8A its close proximity, one residue apart, could explain why the pSer111-effect is not as pronounced for the GAP TBC1D20 as for the Rabin8 interaction (Figure 2A-B and Supplementary Figure 6). Furthermore, we did not detect any influence of Ser111 phosphorylation on the interaction with the effector Mical-1 due to the fact that the Switch II region (α2) of Rab8A is not part of the Mical-1 binding site (Supplementary Figure 6). Arg79 is conserved in Rab1B, and in future studies it will be important to undertake dynamic structural methods such as NMR to study the complexes of pSer111-Rab1B:GEF in comparison to WT Rab1B:GEF to confirm the central role of Arg79 in mediating polar interactions between Rab1B and effectors as well as studies with isotopically labelled Rab8A. Furthermore, it would be interesting to assess whether Arg79 may play a role in mediating specific polar interactions between Rab8A/Rab1B and LRRK2 that is distinct from those of with MST3 or TAK1.

There is great interest in whether the PINK1 signalling pathway converges with other PD gene pathways and whether there are critical signalling nodes mediating neurodegeneration in PD. Few studies have suggested a link between PINK1 and LRRK2. A recent study suggested that LRRK2 mediated kinase activity may inhibit PINK1 mediated mitophagy however, the mechanism was not elaborated [28]. Our data suggests that PINK1-dependent signalling may converge with LRRK2 signalling at Rab8A (Figure 3 and Supplementary Figure 7) and the antagonistic interplay between Ser111 phosphorylation and Thr72 phosphorylation is genetically concordant with how respective mutations in PINK1 and LRRK2 cause Parkinson’s (Figure 6). Endogenous phosphorylation of Thr72 by LRRK2 is low which hampers the ability to assess the interplay of PINK1 and LRRK2 phosphorylation of Rabs. However, recent studies have suggested that effectors including RILPL1 and RILPL2 bind preferentially to LRRK2-phosphorylated Rab Thr72 and this leads to inhibition of primary cilia in the brains of mice harbouring the hyperactive LRRK2 [1441G] mutation [14]. In future work it will be important to test whether cilia inhibition by LRRK2 can be regulated by PINK1 *in vivo* through the comparative analysis of LRRK2 [R1441G]/PINK1 knockout double mutant mice with littermate controls.

**Figure 6.**
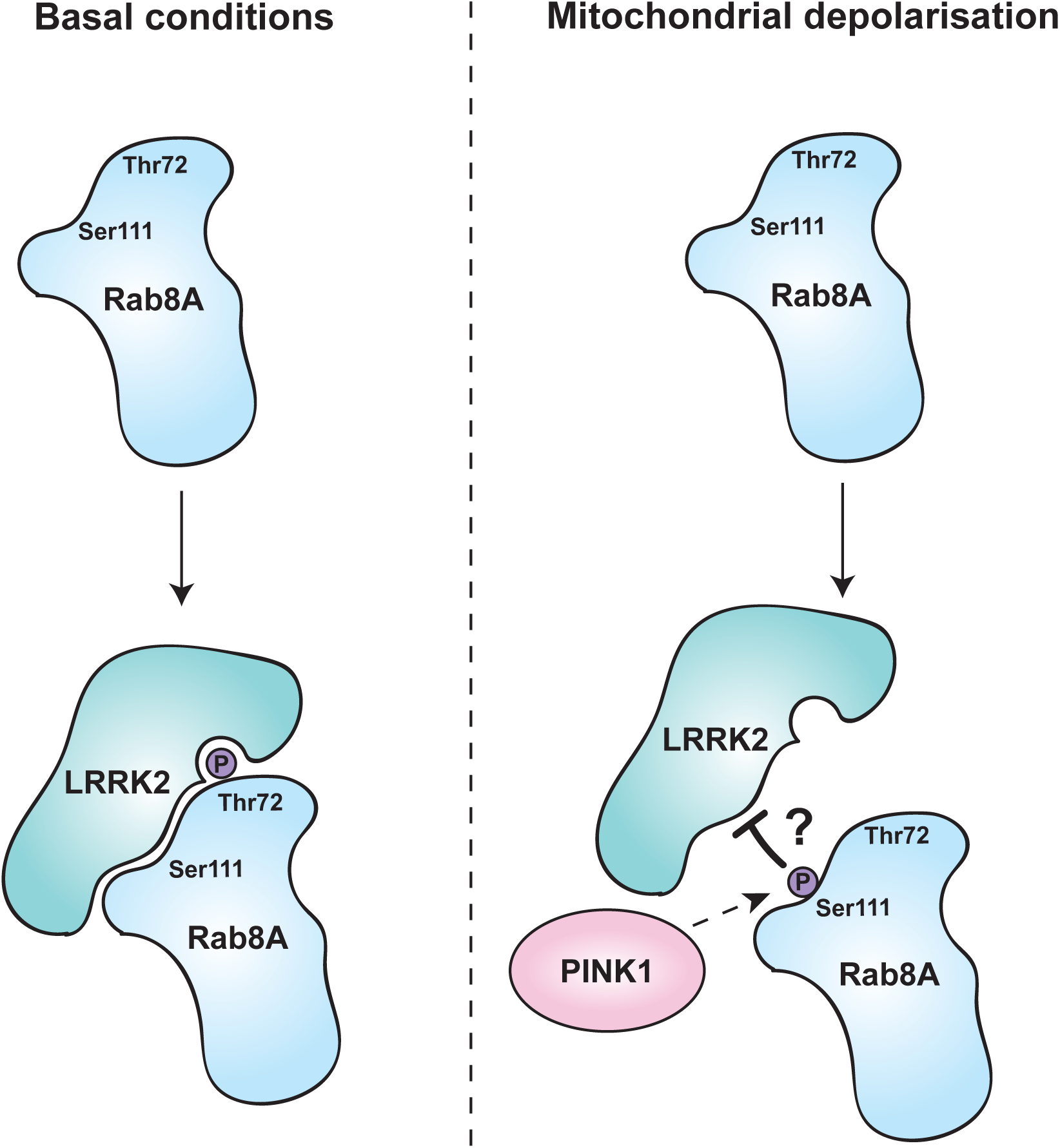
Schematic model of PINK1 and LRRK2 interplay converging on Rab8A. Indirect phosphorylation of Rab8A at Ser111, reduces the ability of LRRK2 to interact and phosphorylate residue Thr72.

Our analysis also revealed that a number of other kinases can regulate Rab phosphorylation including MST3 and TAK1 that are Thr72 directed kinases and MSK1, ZAP70, PAK2, MAP4K5, MAPKAPK3, JNK, PBK, TAK1, and NEK6 whose sites of Rab phosphorylation are unknown. TAK1 has previously been shown to phosphorylate Thr72 in Rab1 and this is important for immunity against Legionella infection [29]. Our studies suggest that the mechanism by which LRRK2 phosphorylates Rab Thr72 is distinct from MST3 and TAK1, and in future it will be important to undertake NMR studies to address this. To date cellular analysis has not suggested an interplay between TAK1 or MST3 with LRRK2 in the regulation mediation of Thr72 of Rabs suggesting that their regulation of Rabs may be distinct from that of LRRK2 biology. However, the discovery of MST3 and TAK1 as Rab8A Thr72 kinases represent new tools to enzymatically purify preparative amounts of Thr72 phosphorylated Rab8A which will accelerate LRRK2 directed research into understanding the consequence of Thr72 phosphorylation on regulation by effectors.

In conclusion our studies have revealed an important role for PINK1-dependent phosphorylation of the SF3 motif of Rabs in the regulation of effector interactions including an antagonistic interplay with LRRK2-mediated Thr72 switch II phosphorylation (Figure 6). In future studies it will be critical to identify the upstream direct kinase for Ser111 since our analysis suggests that activation of the Ser111 kinase may have therapeutic potential to act as a brake on LRRK2-mediated Thr72 phosphorylation.

## Accession codes

Structure factors and coordinates for pSer111-Rab8A:GDP and pSer111-Rab8A:GppNHp were deposited in the RCSB Protein Data Bank (PDB) under the accession codes 6STF and 6STG, respectively.

## Acknowledgements

We express our thanks to Joby Varghese, Robert Gourlay and Renata Soares for mass spectrometry analysis. We thank the laboratory of Prof. Jason W. Chin (Medical Research Council Laboratory of Molecular Biology, Francis Crick Avenue, Cambridge, CB2 0QH) for providing us the plasmids for genetically encoding phospho-serine and the ΔserB-BL21(DE3) *E.coli* cells for expression. We are grateful to the sequencing service (School of Life Sciences, University of Dundee); James Hastie and Hilary McLauchlan and the antibody purification and protein production teams (Division of Signal Transduction Therapy (DSTT), University of Dundee) for excellent technical support. We thank the staff of beamline X06SA at the Paul Scherrer Institute, Swiss Light Source, Villigen, Switzerland, for assistance during data collection. M.M.K.M. is funded by a Wellcome Trust Senior Research Fellowship in Clinical Science (210753/Z/18/Z). This work was supported by the Michael J. Fox Foundation for Parkinson’s disease research, Medical Research Council, and a EMBO Young Investigator Programme Award. K.M. is funded by a Parkinson’s UK PhD studentship. SV, BB, NK, MS, MG, AI acknowledge support by the German Research Foundation (DFG), collaborative research center SFB1035.

## Author Contributions

SV and KM performed most of the experiments. BB and MG performed x-ray crystallography; and NK performed NMR analysis under supervision of MS. Y-CL contributed to *in vitro* kinase screening and validation. RT generated cDNA constructs used in the project. SV, KM, BB, MG, NK, MS, AI and MMKM planned experiments and analysed results. MMKM and AI wrote the paper with contribution from all the authors. MMKM and AI conceived and supervised the project.

---

**Supplementary Figure 1.**
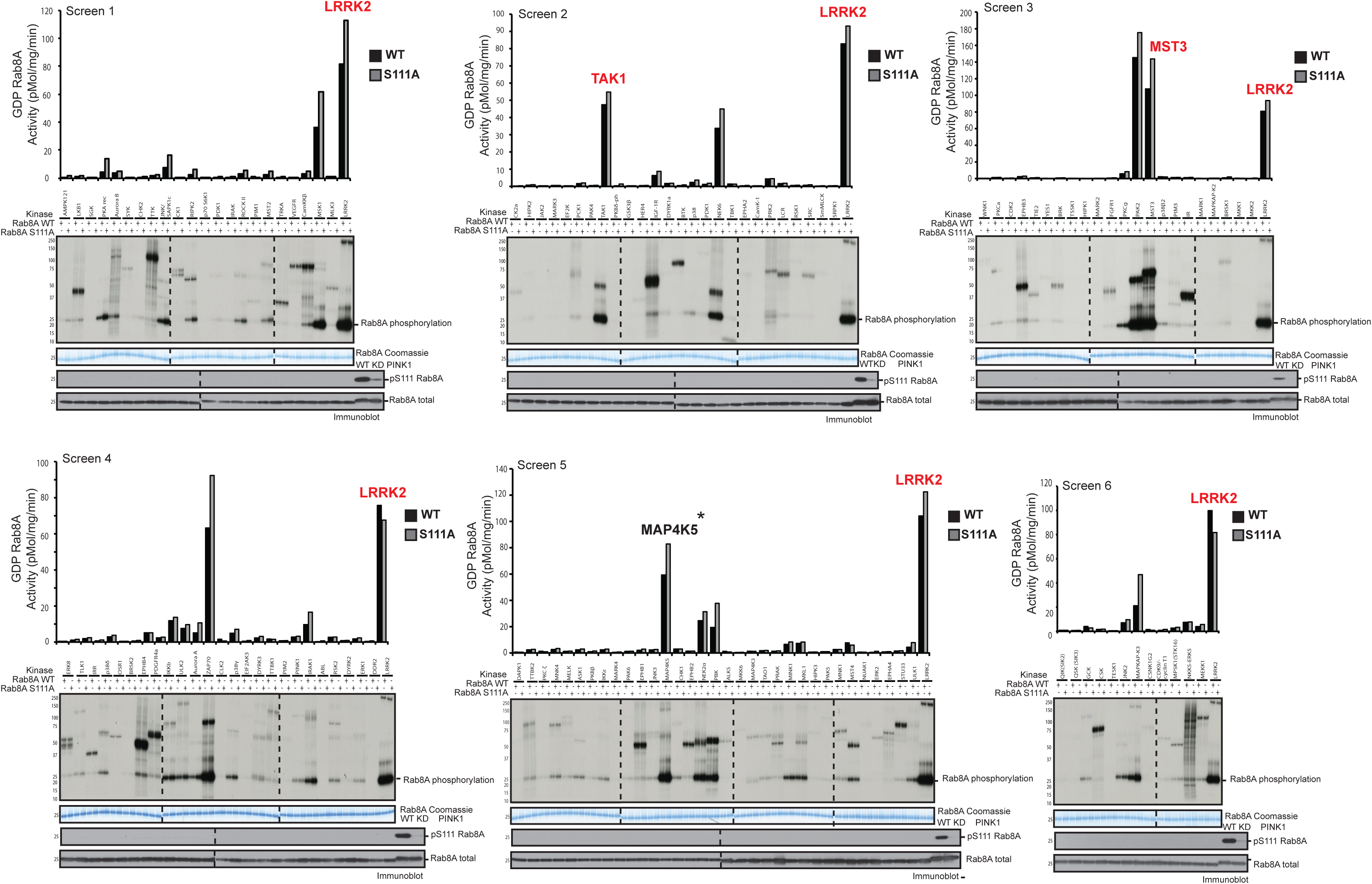
Kinase screen *in vitro* against WT Rab8A:GDP. 100 ng of each protein kinase from the MRC Reagents and Services library was incubated with 2 µg GDP-Rab8A WT or GDP-Rab8A S111A and [γ-^32^P] ATP for 30 min. Samples were subjected to SDS-PAGE with analysis by Coomassie staining, [γ-^32^P] incorporation measured by autoradiography with Cerenkov counting and immunoblotting analysis using the indicated antibodies. Proteins highlighted in red are able to phosphorylate GDP-Rab8A at Thr72.

**Supplementary Figure 2.**
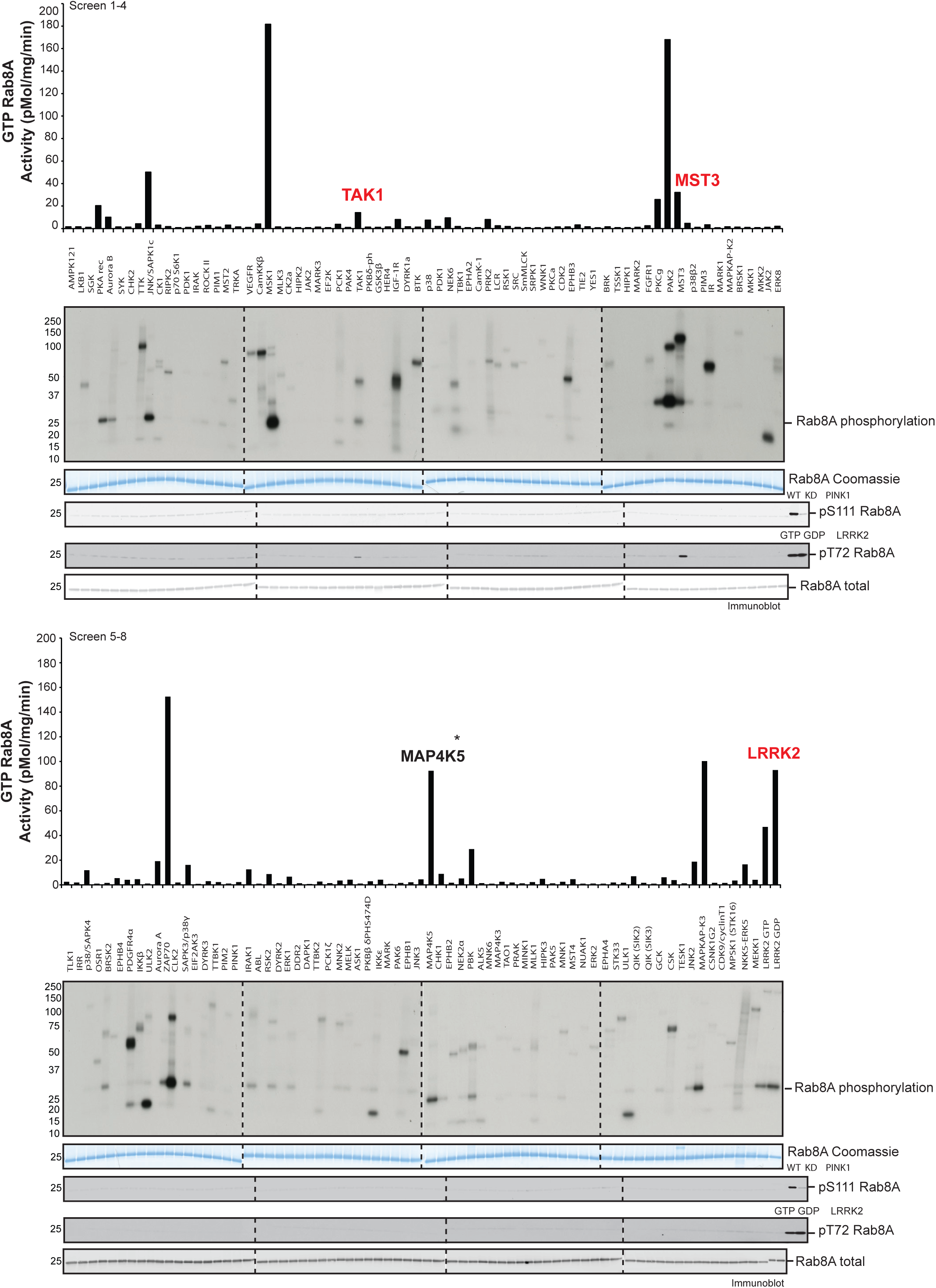
Kinase screen *in vitro* against WT Rab8A: GppNHp. 100 ng of each protein kinase from the MRC Reagents and Services library was incubated with 2 µg WT Rab8A:GppNHp and [γ-^32^P] ATP for 30 min. Samples were subjected to SDS-PAGE with analysis by Coomassie staining, [γ-^32^P] incorporation measured by autoradiography with Cerenkov counting and immunoblotting analysis using the indicated antibodies. Proteins highlighted in red are able to phosphorylate WT Rab8A:GppNHp at Thr72.

**Supplementary Figure 3.**
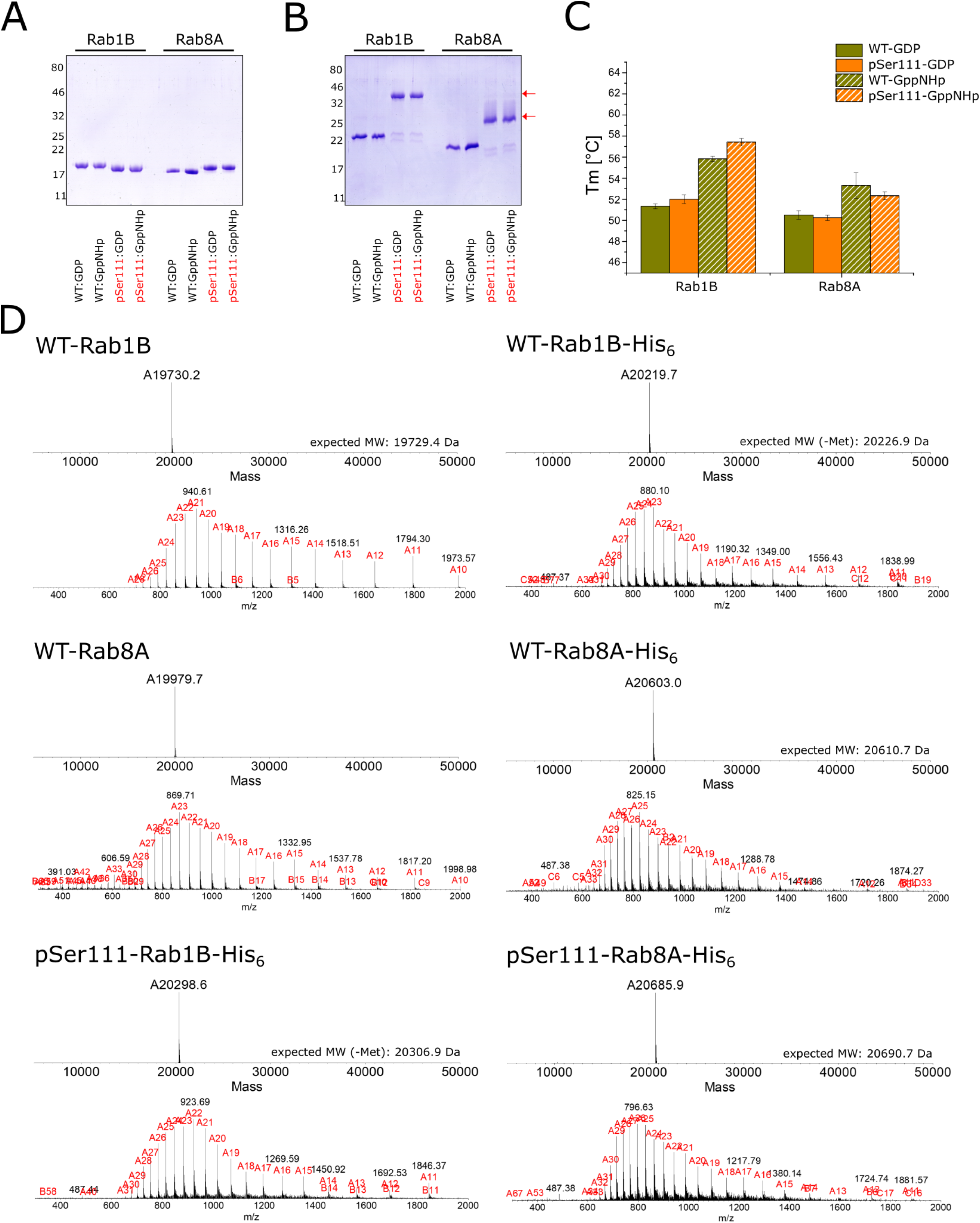
Expression and purification of Ser111-phosphorylated Rab1B and 8A GTPases. **A-B.** Analysis of the purity of WT-Rab1B/8a-His_6_ and pSer111-Rab1B/8A-His_6_ proteins by 12% SDS-PAGE (A). Detection of phosphorylated amino acid residues in WT-Rab1B/8a-His_6_ and pSer111-Rab1B/8a-His_6_ proteins by Phos-tag™ acrylamide SDS-PAGE. The migration of phosphorylated Rab proteins is retarded (highlighted with red arrows) due to the phos-tag™ compound (B). **C:** Melting points (Tm) of the WT-Rab1B/8a and the pSer111-Rab1B/8a-His_6_ proteins determined by a SYPRO® orange-based thermal shift assay. **D:** LC-MS analysis of the WT- and pSer111-Rab proteins used in this study. Note that the N-terminal methionine of Rab1B is usually cleaved off during expression.

**Supplementary Figure 4.**
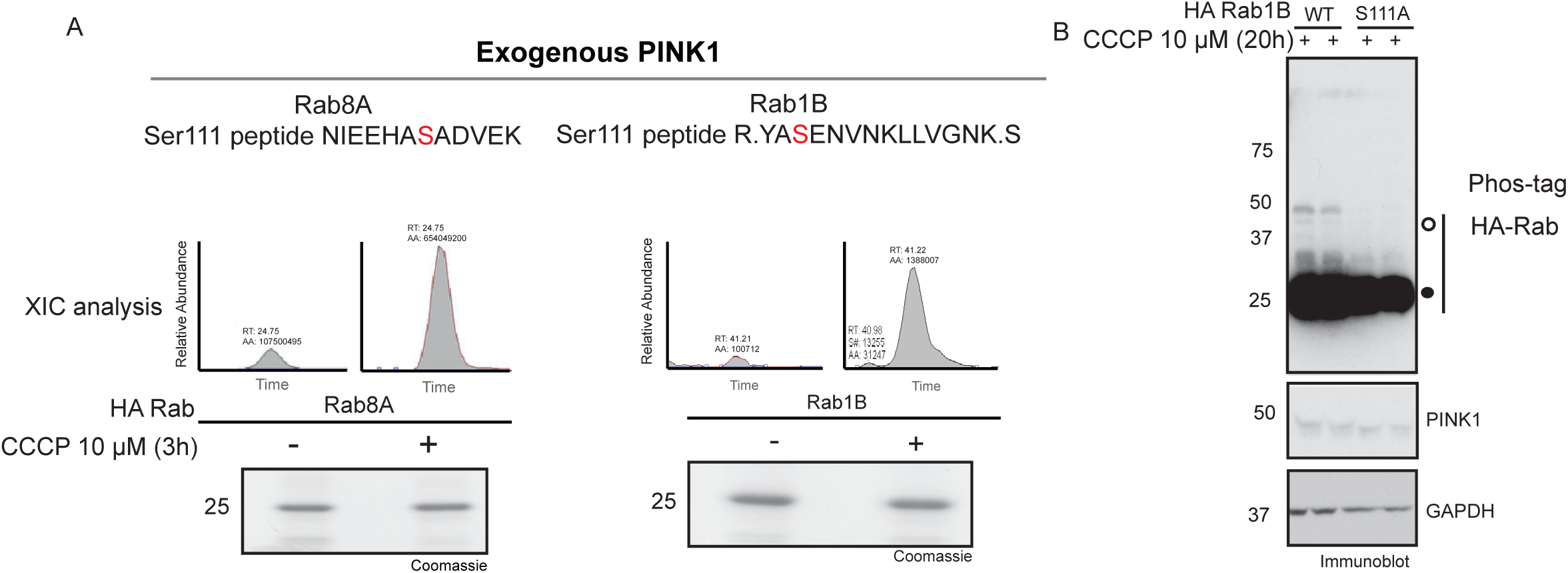
Rab1B is indirectly phosphorylated by PINK1 at Ser111. **A.** HEK293 cells stably expressing 3XFlag PINK1 WT were transfected with either Rab8A WT or Rab1B WT and stimulated with 10 µM CCCP for 3 h. 10 mg of whole cell lysate was immunoprecipitated with anti-HA agarose, resolved by SDS-PAGE and stained with colloidal Coomassie. Displayed bands were excised, followed by trypsin digestion and subjected to high performance liquid chromatography with tandem mass spectrometry (LC-MS-MS) on an LQT-Oribitrap mass spectrometer. XIC’s display the absolute area (AA) of each Ser111 phosphopeptide of interest, with Y-axis corresponding to phosphopeptide signal intensity and x-axis to the retention time (RT). **B.** HeLa cells were transiently transfected with either HA-Rab1B WT or S111A constructs, followed by 10 μM CCCP or DMSO treatment for 20 h. Samples were subjected to Phos-tag analysis.

**Supplementary Figure 5.**
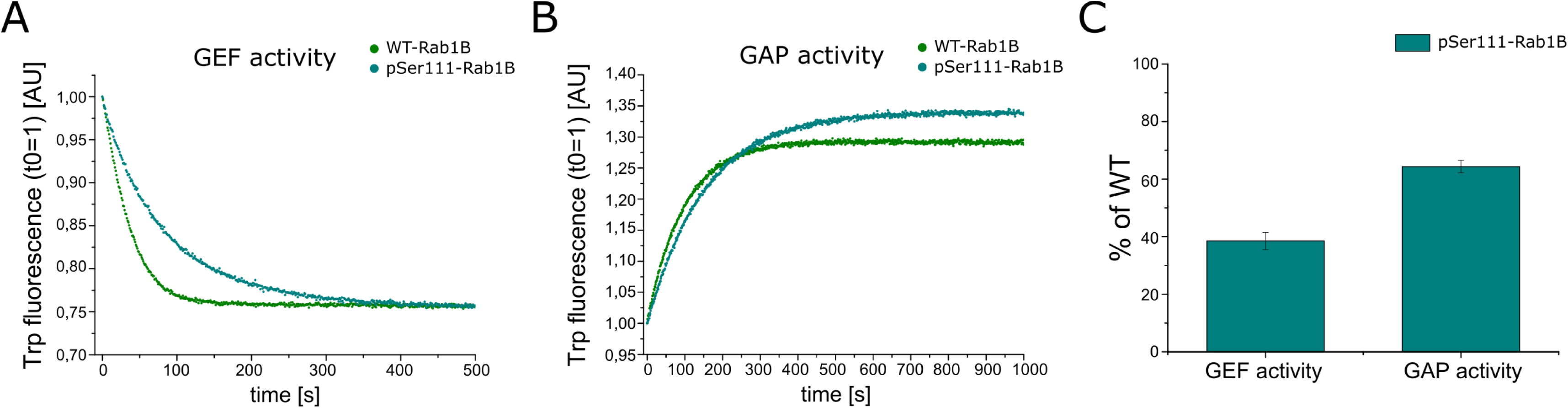
Rab1B Ser111 phosphorylation impairs GEF and GAP interactions. **A, B.** (A) The GDP-GTP exchange catalyzed by the GEF DrrA, and (B) the GTP hydrolysis catalyzed by the GAP TBC1D20 (B) were monitored for WT-Rab1B (green) and pSer111-Rab1B-His_6_ (teal) proteins via intrinsic tryptophan (Trp) fluorescence. The shown graphs are representative for three independent experiments. C) The nucleotide exchange (GEF activity) and GTP hydrolysis rates (GAP activity) were determined using a single exponential fit, and are represented as % of WT. All data (n=3) are represented as mean ± S.D.

**Supplementary Figure 6.**
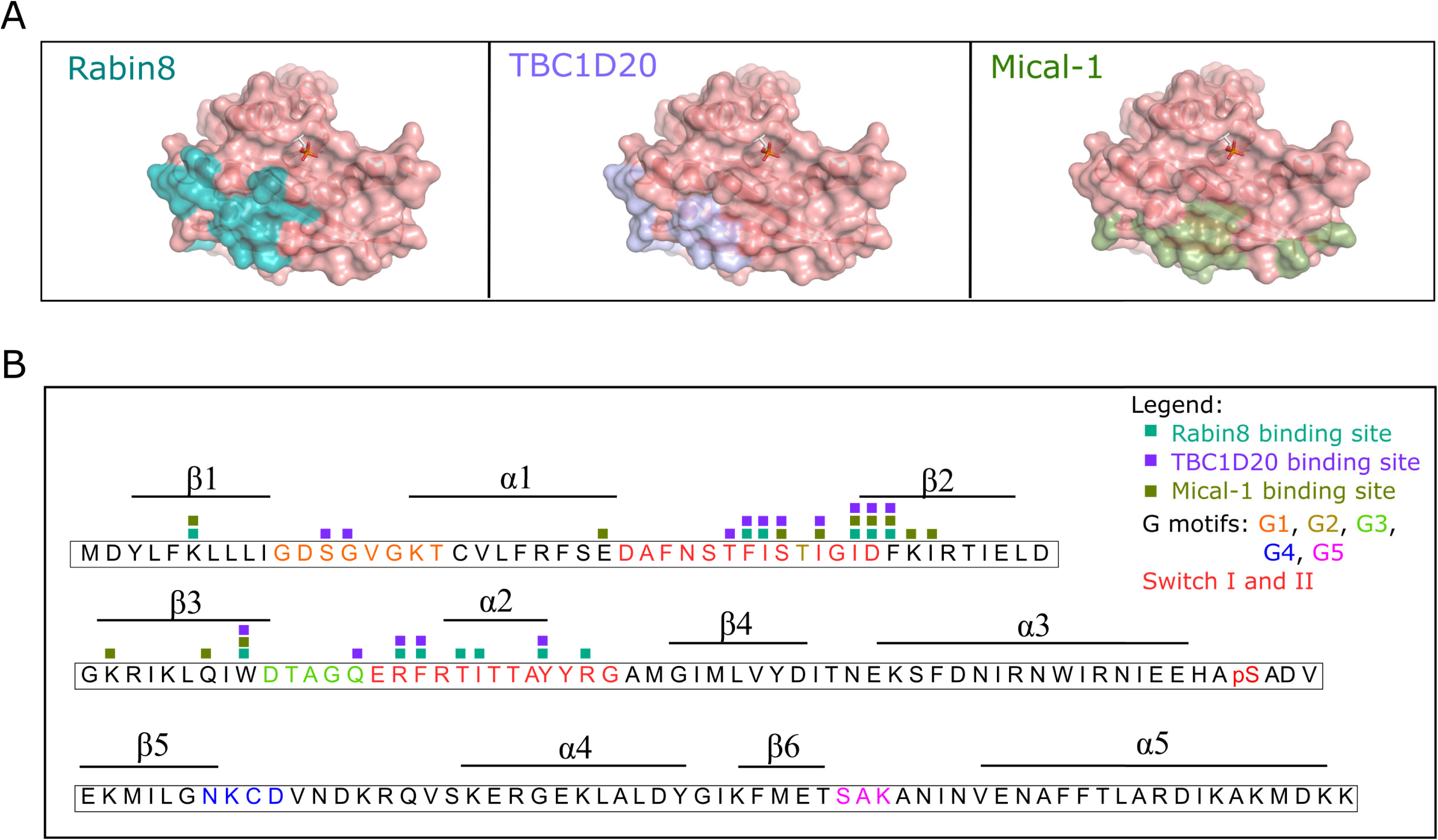
Binding sites of Rab8A regulators and effectors. **A.** A depiction of the binding sites of the GEF Rabin8 [21] (teal), the GAP TBC1D20 [40, 41] (purple), and the effector Mical-1 [18] (olive green) on the surface of pSer111-Rab8A:GppNHp (salmon). **B.** Schematic representation of the pSer111-Rab8A amino acid sequence and secondary structure highlighting the binding sites of Rabin8 (teal), TBC1D20 (purple) and Mical-1 (olive green).

**Supplementary Figure 7.**
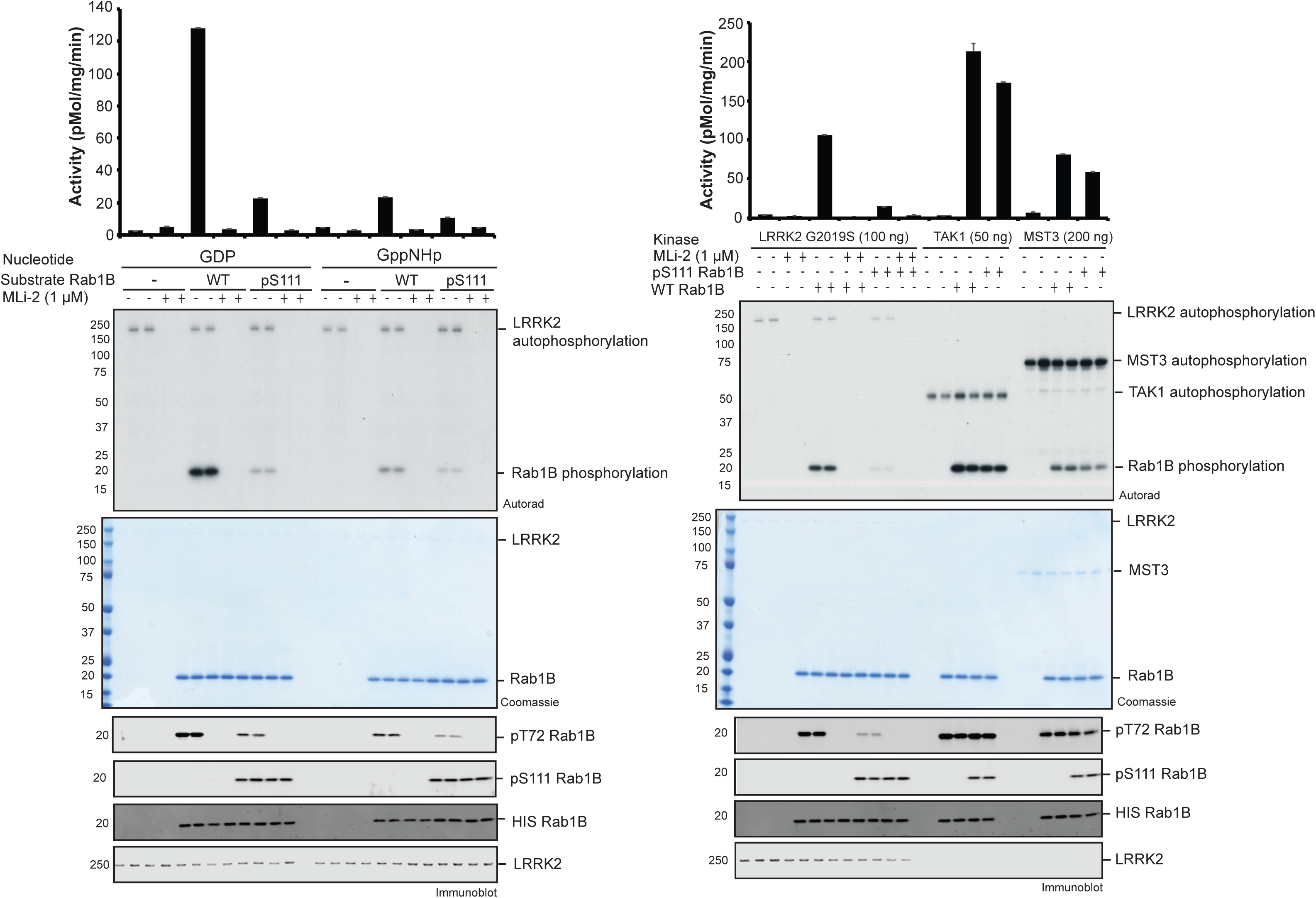
Rab1B Ser111 phosphorylation disrupts LRRK2-mediated Thr72 phosphorylation *in vitro*. **A.** 100 ng of recombinant LRRK2 [G2019S] protein was incubated with [γ-^32^P] and 2 µg of WT or pSer114 Rab1B, in either the GDP or GTP (GppNHp) bound conformation, in the absence or presence of the LRRK2 inhibitor, MLi-2 for 30 min. Samples were subjected to SDS-PAGE, and analysed by either Coomassie staining, [γ-^32^P] incorporation measured by autoradiography with Cerenkov counting (top panel) and immunoblot analysis, using the indicated antibodies (lower panel). Results are means +/- S.E.M. (*n*=3). **B.** Thr72 specific kinases were assessed for phosphorylation of GDP bound WT or pSer114 Rab1B. 100 ng LRRK2 G2019S, 50 ng TAK1 or 200 ng MST3 were incubated with 2 µg substrate using identical conditions and analysis as in panel A.

**Supplementary Figure 8.**
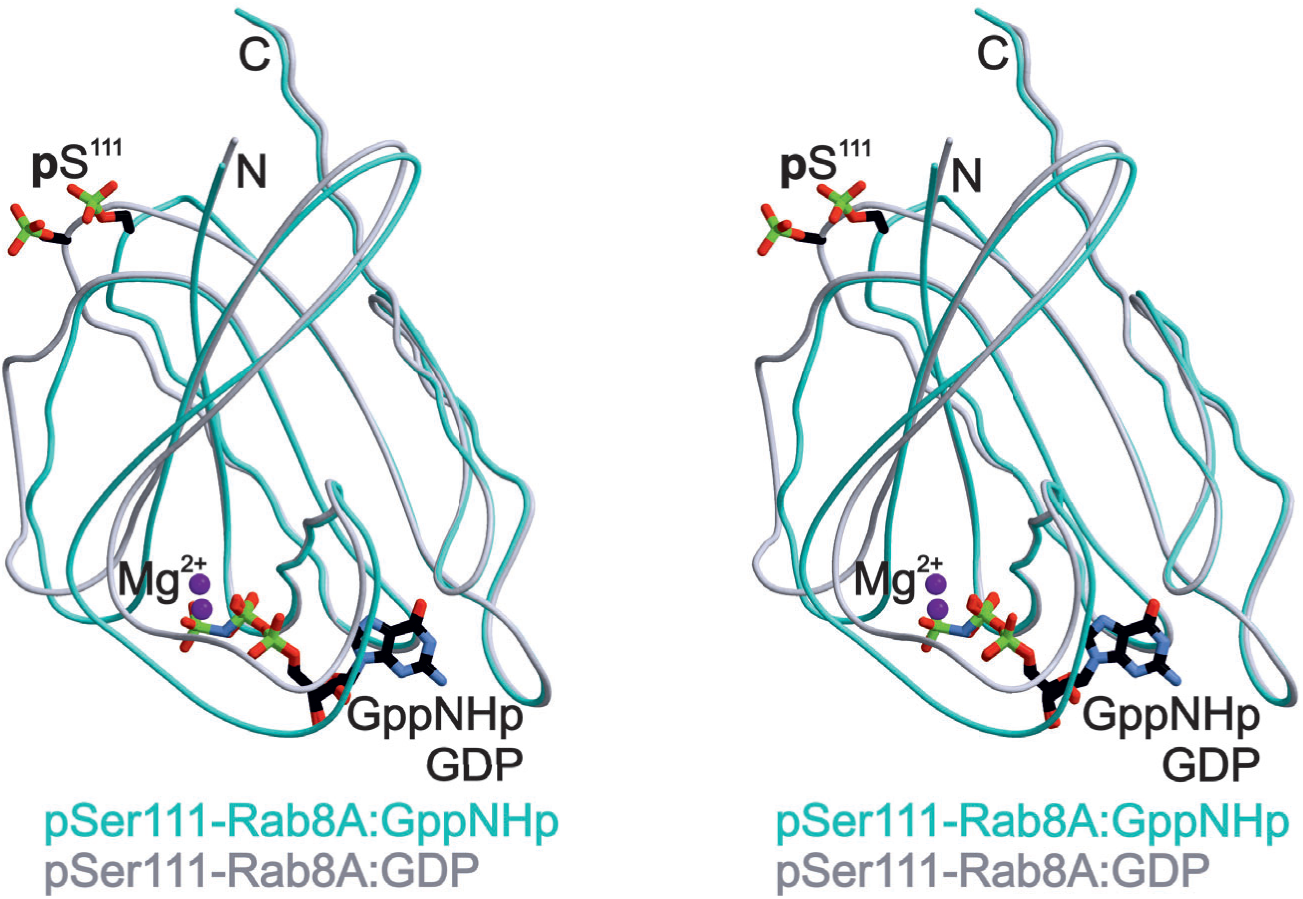
Superposition of x-ray structures of Rab8A phosphorylated at Ser111. Stereo view of the structural superposition of pSer111-Rab8A:GppNHp with pSer111-Rab8A:GDP.

**Supplementary Figure 9.**
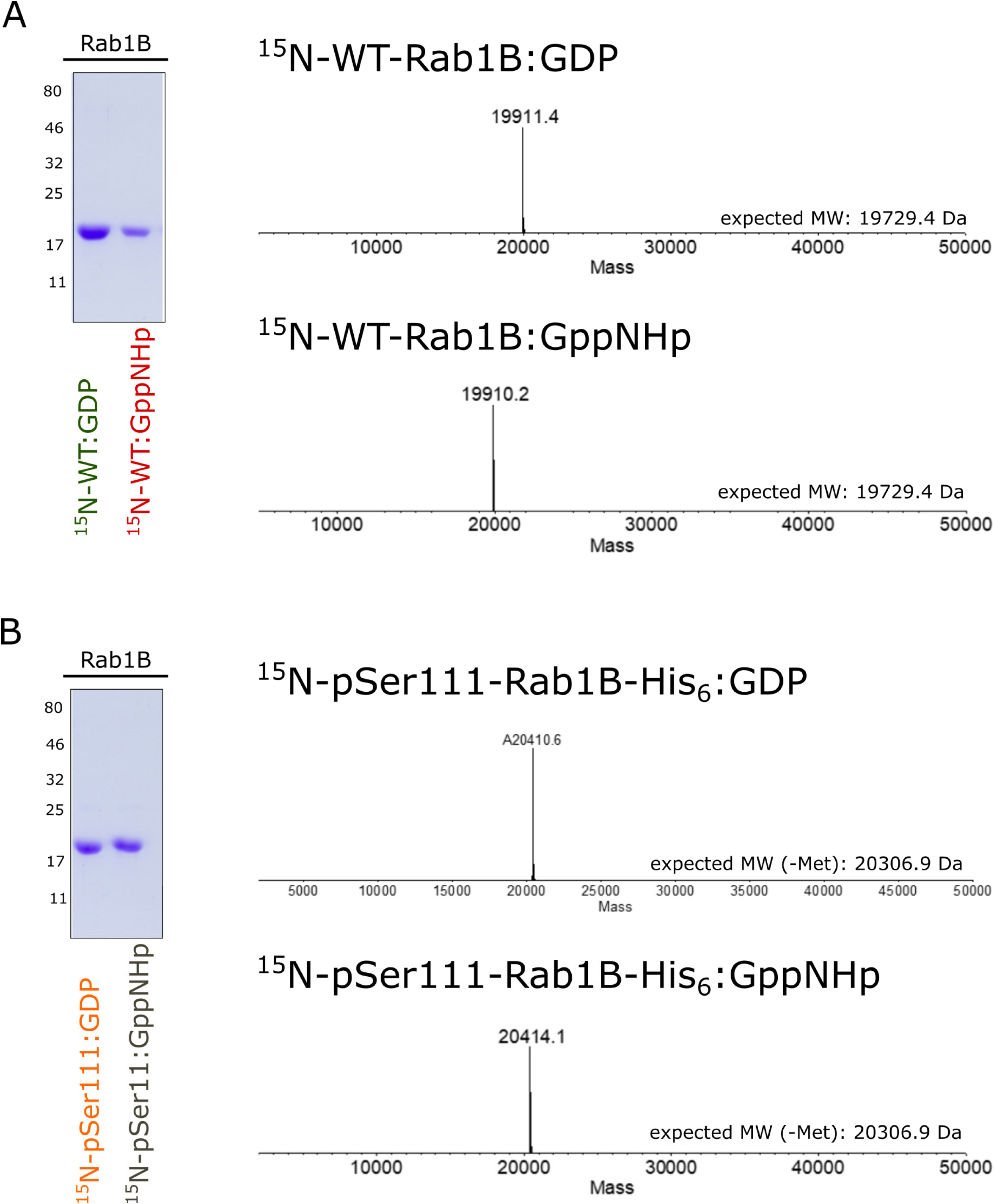
Analytical characterization of ^15^N-labelled WT- and pSer111-Rab1B proteins. **A.** Analysis of ^15^N-labelled WT-Rab1B:GDP/GppNHp proteins by LC-MS and SDS-PAGE (12%). **B.** Analysis of the ^15^N-labelled pSer111-Rab1B:GDP/GppNHp proteins by LC-MS and SDS-PAGE (12%). Note that the N-terminal methionine of Rab1B is usually cleaved off during expression.

**Supplementary Figure 10.**
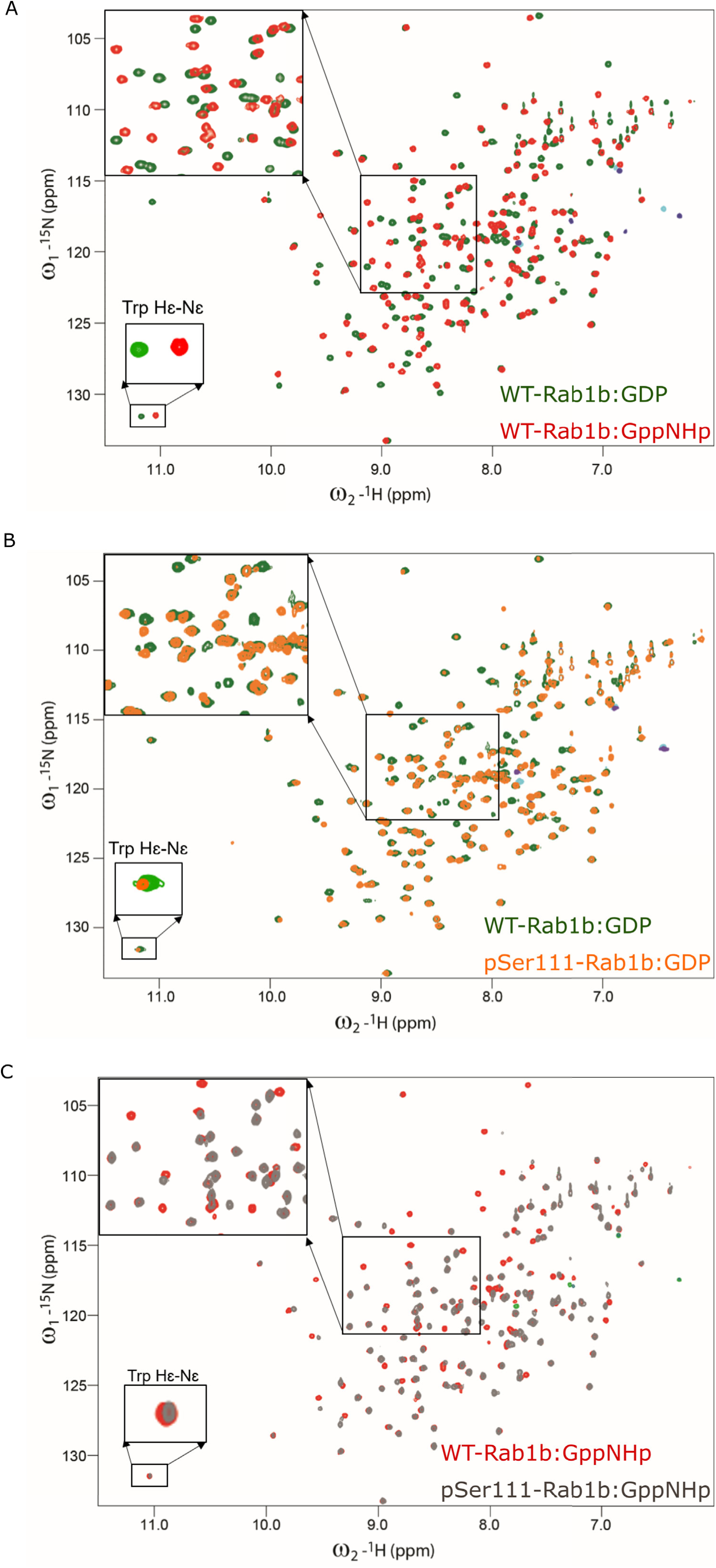
^1^H-^15^N HSQC solution NMR spectra of WT- and pSer111-Rab1B spectra in the inactive and active state. Figure insets show the zoom region for central and Trp Hɛ-Nɛ. **A.** Overlay spectra of WT-Rab1B:GDP (green) and WT-Rab1B:GppNH (red). Major changes between inactive vs active conformations are shown via overlapping of the WT-Rab1B:GDP and WT-Rab1B:GppNH spectra. **B.** Inactive conformations of unmodified and phosphorylated Rab1B. Overlay spectra shows WT-Rab1B:GDP (green) and pSer111-Rab1B:GDP in orange. Inset shows the central zoom area of the spectra. **C.** Active conformation of unmodified and phosphorylated Rab1B. Overlay spectra of WT-Rab1B:GppNH (red) and pSer111-Rab1B:GppNH (gray).

**Supplementary Figure 11.**
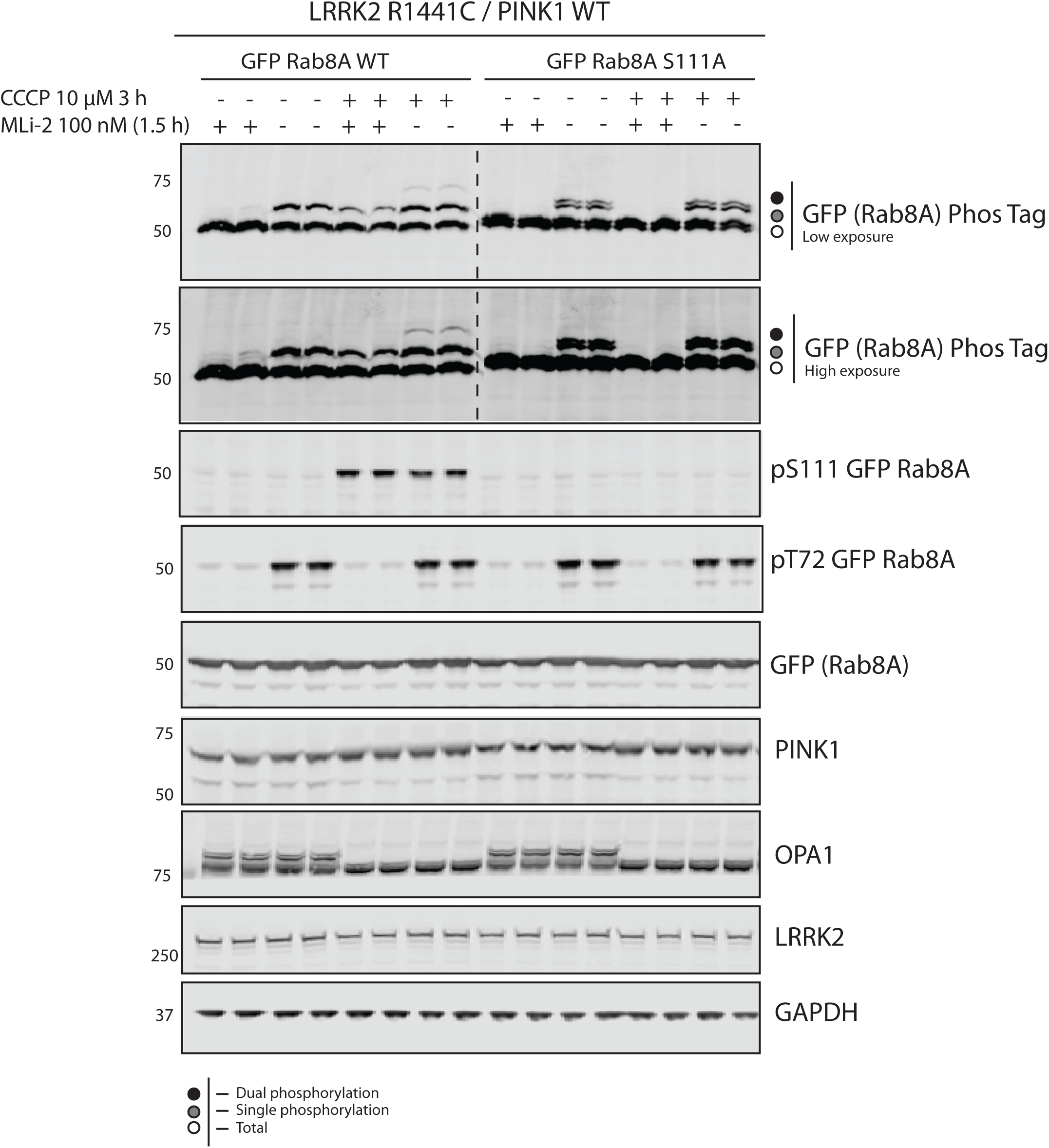
PINK1 and LRRK2 signalling converge on Rab8A in cells.: loading controls and mitochondrial depolarisation readouts. HEK293 Flp-In TREx cells stably expressing PINK1-3XFLAG were co-transfected with 9 μg of LRRK2 R1441G and 3 μg of GFP Rab8A WT or S111A. Cells were treated with DMSO, 100 nM MLi-2 (1.5 h), 10 μM CCCP (3 h) or a combination of each. Cells were lysed and underwent immunoblotting with Phos-tag analysis utilising total antibodies for experimental controls and phospho-specific antibodies as experimental read-outs. Anti-GAPDH and Anti-LRRK2 antibodies confirmed equal loading and LRRK2 expression. Anti-OPA1 antibody confirmed mitochondrial depolarisation across samples. Immunoblots were visualised using the LICOR Odyssey imaging system.

## References

1. Valente, E.M., et al., Hereditary early-onset Parkinson’s disease caused by mutations in PINK1. Science, 2004. 304(5674): p. 1158-60.

2. Kondapalli, C., et al., PINK1 is activated by mitochondrial membrane potential depolarization and stimulates Parkin E3 ligase activity by phosphorylating Serine 65. Open Biol, 2012. 2(5): p. 120080.

3. Shiba-Fukushima, K., et al., PINK1-mediated phosphorylation of the Parkin ubiquitin-like domain primes mitochondrial translocation of Parkin and regulates mitophagy. Sci Rep, 2012. 2: p. 1002.

4. Kane, L.A., et al., PINK1 phosphorylates ubiquitin to activate Parkin E3 ubiquitin ligase activity. J Cell Biol, 2014. 205(2): p. 143–53.

5. Kazlauskaite, A., et al., Parkin is activated by PINK1-dependent phosphorylation of ubiquitin at Ser65. Biochem J, 2014. 460(1): p. 127–39.

6. Koyano, F., et al., Ubiquitin is phosphorylated by PINK1 to activate parkin. Nature, 2014. 510(7503): p. 162-6.

7. Ordureau, A., et al., Quantitative proteomics reveal a feedforward mechanism for mitochondrial PARKIN translocation and ubiquitin chain synthesis. Mol Cell, 2014. 56(3): p. 360–75.

8. Lai, Y.C., et al., Phosphoproteomic screening identifies Rab GTPases as novel downstream targets of PINK1. EMBO J, 2015. 34(22): p. 2840–61.

9. Steger, M., et al., Systematic proteomic analysis of LRRK2-mediated Rab GTPase phosphorylation establishes a connection to ciliogenesis. Elife, 2017. 6.

10. Steger, M., et al., Phosphoproteomics reveals that Parkinson’s disease kinase LRRK2 regulates a subset of Rab GTPases. Elife, 2016. 5.

11. Mir, R., et al., The Parkinson’s disease VPS35[D620N] mutation enhances LRRK2-mediated Rab protein phosphorylation in mouse and human. Biochem J, 2018. 475(11): p. 1861–1883.

12. Pylypenko, O., et al., Rab GTPases and their interacting protein partners: Structural insights into Rab functional diversity. Small GTPases, 2018. 9(1-2): p. 22–48.

13. Muller, M.P. and R.S. Goody, Molecular control of Rab activity by GEFs, GAPs and GDI. Small GTPases, 2018. 9(1-2): p. 5–21.

14. Dhekne, H.S., et al., A pathway for Parkinson’s Disease LRRK2 kinase to block primary cilia and Sonic hedgehog signaling in the brain. Elife, 2018. 7.

15. Rogerson, D.T., et al., Efficient genetic encoding of phosphoserine and its nonhydrolyzable analog. Nat Chem Biol, 2015. 11(7): p. 496–503.

16. Muller, M.P., et al., Characterization of enzymes from Legionella pneumophila involved in reversible adenylylation of Rab1 protein. J Biol Chem, 2012. 287(42): p. 35036–46.

17. Vanoni, M.A., Structure-function studies of MICAL, the unusual multidomain flavoenzyme involved in actin cytoskeleton dynamics. Arch Biochem Biophys, 2017. 632: p. 118–141.

18. Rai, A., et al., bMERB domains are bivalent Rab8 family effectors evolved by gene duplication. Elife, 2016. 5.

19. Schlichting, I., et al., Time-resolved X-ray crystallographic study of the conformational change in Ha-Ras p21 protein on GTP hydrolysis. Nature, 1990. 345(6273): p. 309-15.

20. Stenmark, H. and V.M. Olkkonen, The Rab GTPase family. Genome Biol, 2001. 2(5): p. REVIEWS3007.

21. Guo, Z., et al., Intermediates in the guanine nucleotide exchange reaction of Rab8 protein catalyzed by guanine nucleotide exchange factors Rabin8 and GRAB. J Biol Chem, 2013. 288(45): p. 32466–74.

22. Ostermeier, C. and A.T. Brunger, Structural basis of Rab effector specificity: crystal structure of the small G protein Rab3A complexed with the effector domain of rabphilin-3A. Cell, 1999. 96(3): p. 363–74.

23. Pereira-Leal, J.B. and M.C. Seabra, The mammalian Rab family of small GTPases: definition of family and subfamily sequence motifs suggests a mechanism for functional specificity in the Ras superfamily. J Mol Biol, 2000. 301(4): p. 1077–87.

24. Hutagalung, A.H. and P.J. Novick, Role of Rab GTPases in membrane traffic and cell physiology. Physiol Rev, 2011. 91(1): p. 119–49.

25. Pfeffer, S. and D. Aivazian, Targeting Rab GTPases to distinct membrane compartments. Nat Rev Mol Cell Biol, 2004. 5(11): p. 886–96.

26. Wu, Y.W., et al., Membrane targeting mechanism of Rab GTPases elucidated by semisynthetic protein probes. Nat Chem Biol, 2010. 6(7): p. 534–40.

27. Pourjafar-Dehkordi, D., et al., Phosphorylation of Ser111 in Rab8a Modulates Rabin8-Dependent Activation by Perturbation of Side Chain Interaction Networks. Biochemistry, 2019. 58(33): p. 3546–3554.

28. Bonello, F., et al., LRRK2 impairs PINK1/Parkin-dependent mitophagy via its kinase activity: pathologic insights into Parkinson’s disease. Hum Mol Genet, 2019. 28(10): p. 1645–1660.

29. Levin, R.S., et al., Innate immunity kinase TAK1 phosphorylates Rab1 on a hotspot for posttranslational modifications by host and pathogen. Proc Natl Acad Sci U S A, 2016. 113(33): p. E4776–83.

30. Fell, M.J., et al., MLi-2, a Potent, Selective, and Centrally Active Compound for Exploring the Therapeutic Potential and Safety of LRRK2 Kinase Inhibition. J Pharmacol Exp Ther, 2015. 355(3): p. 397-409.

31. Shaw Stewart, P.D., Kolek, S.A., Briggs, R.A., Chayen, N.E, Baldock, P.F.M., Random Microseeding: A Theoretical and Practical Exlporation of Seed Stability and Seeding Techniques for Successful Protein Crystallisation. Cryst. Growth Des, 2011. 11: p. 3432–3441.

32. Kabsch, W., Integration, scaling, space-group assignment and post-refinement. Acta Crystallogr D Biol Crystallogr, 2010. 66(Pt 2): p. 133–44.

33. McCoy, A.J., et al., Phaser crystallographic software. J Appl Crystallogr, 2007. 40(Pt 4): p. 658–674.

34. Emsley, P. and K. Cowtan, Coot: model-building tools for molecular graphics. Acta Crystallogr D Biol Crystallogr, 2004. 60(Pt 12 Pt 1): p. 2126-32.

35. Adams, P.D., et al., PHENIX: a comprehensive Python-based system for macromolecular structure solution. Acta Crystallogr D Biol Crystallogr, 2010. 66(Pt 2): p. 213–21.

36. Langer, G., et al., Automated macromolecular model building for X-ray crystallography using ARP/wARP version 7. Nat Protoc, 2008. 3(7): p. 1171–9.

37. Chen, V.B., et al., MolProbity: all-atom structure validation for macromolecular crystallography. Acta Crystallogr D Biol Crystallogr, 2010. 66(Pt 1): p. 12–21.

38. Vranken, W.F., et al., The CCPN data model for NMR spectroscopy: development of a software pipeline. Proteins, 2005. 59(4): p. 687–96.

39. Reed, S.E., et al., Transfection of mammalian cells using linear polyethylenimine is a simple and effective means of producing recombinant adeno-associated virus vectors. J Virol Methods, 2006. 138(1-2): p. 85–98.

40. Pan, X., et al., TBC-domain GAPs for Rab GTPases accelerate GTP hydrolysis by a dual-finger mechanism. Nature, 2006. 442(7100): p. 303-6.

41. Gavriljuk, K., et al., Catalytic mechanism of a mammalian Rab.RabGAP complex in atomic detail. Proc Natl Acad Sci U S A, 2012. 109(52): p. 21348–53.

